# Nature-Inspired Nanoparticle Adiposomes Enable Targeted Delivery of Hydrophobic Drug for Anti-Cancer Treatment

**DOI:** 10.64898/2026.06.01.729180

**Authors:** Bin Pan, Zhen Cao, Zemin Li, Gaoxin Zhang, Chang Zhou, Zelun Zhi, Yanqiu Zhu, Qi Zhang, Shasha Lu, Shuyan Zhang, Yuhan Zhao, Bing Yan, Xintong Li, Kevin Xiaohu Liu, Pingsheng Liu

## Abstract

The predominance of hydrophobic molecules in drug pipelines poses a fundamental delivery challenge in the aqueous physiological environment. Inspired by the endogenous lipid-transport machinery of lipid droplets and lipoproteins, adiposome was developed as a biocompatible platform for hydrophobic drug delivery. Here, docetaxel (DTX) was selected as a model hydrophobic chemotherapeutic agent to evaluate the therapeutic versatility of adiposome. Non-targeted DTX-loaded adiposomes (DTX-Ad) reduced systemic toxicity by avoiding the allergenic excipients Tween 80 and ethanol used in commercial DTX formulations. The cytotoxic mechanism and intracellular responses of DTX-Ad were comprehensively characterized. To boost antitumor efficacy, three tumor-targeting DTX-Ad formulations were subsequently engineered. Lung-targeting DTX-adiposomes (Lung@DTX-Ad), mediated by AAM-B_1-28_-CRGDK fusion peptide/NRP1 axis, enhanced antitumor efficacy in a lung metastasis model. BCMA-targeting DTX-adiposomes (BCMA@DTX-Ad), conferred by biotin-avidin-mediated antibody conjugation, achieved sustained suppression in a multiple myeloma model. Liver-targeting DTX-adiposomes (Liver@DTX-Ad), enabled by the ApoE/LDLR axis, outperformed both commercial DTX injection and sorafenib in hepatocellular carcinoma models. Taken together, these results demonstrate adiposome as a promising and broadly applicable drug delivery platform, with considerable potential for clinical translation across diverse malignancies.

## Introduction

Detailed knowledge of molecular mechanisms of varieties of human diseases has been identified and accumulated since the completion of the Human Genome Project^[1]^ as well as the application of artificial intelligence (AI)^[2]^, which greatly facilitates the development of new therapeutic intracellular targets of the diseases. The newly developed small molecules are more hydrophobic^[3]^ because many compounds must cross the plasma membrane that consists of bilayer of hydrophobic fatty acids^[4]^. Therefore, it is estimated that approximately 40% of drugs in the market and 90% of molecules candidates in the development pipelines are poorly water-soluble^[5]^. This raises a dilemma of delivery of the hydrophobic molecules in the hydrophilic environment of blood.

This hydrophobicity challenge is deeply rooted not only in the drug delivery but also in the development of drug targets. Previously, G protein-coupled receptors (GPCRs) had been the most successful drug targets, with about 34-40% of FDA-approved drugs targeting them^[6; 7]^. Owing to the features of GPCR structure as seven transmembrane receptors, these drugs mainly act at extracellular or transmembrane binding sites. Therefore, drugs targeting GPCRs are mainly hydrophilic molecules that can be delivered in blood system. In contrast, modern drug discovery programs have mostly shifted to intracellular targets^[8]^. The plasma membrane is the first barrier that prevents direct translocation of chemical entities. To cross the cell membrane, hydrophobic small molecule drugs with high permeability is necessarily developed. However, these drugs encounter many challenges in water solubility, drug resistance, subcellular targeting, and controlled release^[9]^.

To facilitate delivery of hydrophobic drug molecules, many approaches have been developed including solvents, vehicles, and many types of nanoparticles^[5]^. With different specific abilities, these therapeutic delivery tools advance the usage of many hydrophobic drugs. Nanoparticles have been widely developed and employed for the encapsulation and delivery of hydrophobic drugs, including liposomes, solid lipid nanoparticles (SLNs), and polymeric micelles^[10]^. However, their intrinsic properties also limit drug-delivery performance, such as low stability, limited drug-loading capacity, and lack of ability to deliver the drug to target cells. Hydrophobic drugs can only be incorporated within the hydrophobic region of the phospholipid bilayer of liposomes, resulting in membrane perturbation and drug leakage^[11]^. SLNs are made up of crystal solid lipids surrounded by surfactants at room temperature. The preparation process of SLNs is prone to disturbing the drug stability due to the hot homogenization and cooling crystallization^[12]^. Non-natural polymers and PEGylated lipids may induce immune responses and alter drug pharmacokinetics/pharmacodynamics^[13]^. Therefore, it is desirable to develop potent delivery systems with biocompatibility and precise targeting for achieving effective intracellular delivery of hydrophobic entities.

Intriguingly, endogenous biological systems have evolved dedicated machinery for the efficient transport of hydrophobic molecules in aqueous environments. In the human body, most hydrophobic molecules are transported in the blood through the lipoprotein system. Lipoproteins (LPs) are endogenous particles composed of a neutral lipid core surrounded by a monolayer phospholipid membrane and apolipoproteins^[14; 15]^. Lipid droplets (LDs) share a similar architecture and store neutral lipids or hydrophobic molecules inside cells. Inspired by the LPs and LDs, which are natural nanoparticles for reserving and transporting hydrophobic substances to maintain homeostasis, a novel nanoparticle termed the adiposome is developed by our group^[16]^. Unlike liposomes and exosomes, which comprised of an aqueous core surrounded by a phospholipid bilayer membrane^[17]^, adiposome contains a neutral lipid core wrapped with a monolayer-phospholipid membrane. Adiposome is a powerful *in vitro* system to study the functions of LDs and LPs, for example, LD fusion^[18; 19]^, LD-associated protein binding affinity^[20-22]^, lipase enzyme activity^[23; 24]^, and LP functions^[25; 26]^. More importantly, adiposome is an ideal nanoparticle for the delivery of water-insoluble drugs due to its large neutral lipid core, good biocompatibility, and tumor-targeting modification.

To demonstrate the translational potential of adiposome as a drug delivery platform, we selected docetaxel (DTX) as a model hydrophobic chemotherapeutic agent. DTX is one of the most effective chemotherapeutic drugs with wide application in different types of cancers, including breast cancer, ovarian cancer, gastric cancer, head and neck cancer^[27-29]^. DTX binds to β-tubulin in microtubules and inhibits their depolymerization, leading to cell death by cell cycle arrest^[30]^. DTX shows a higher cytotoxicity than paclitaxel with a similar structure and antitumor mechanism^[31]^. However, because of poor-water solubility, intolerable side effects and non-tumor targeting, the clinical efficacy of DTX is substantially limited^[32]^. Hence, nature-inspired adiposome is an efficient delivery system that is engineered to address the aforementioned drawbacks.

Here, we first generated non-targeted DTX-loaded adiposomes (DTX-Ad) and comprehensively evaluated its intracellular responses by a variety of analytical methods. Compared with commercial docetaxel injection (DTX-Inj), DTX-Ad diminished systemic toxicities due to eliminating allergenic excipients, Tween 80 and ethanol. To boost its antitumor efficacy, we creatively engineered various targeted DTX-Ad formulations. First, lung-targeting DTX-adiposomes (Lung@DTX-Ad) were designed and showed stronger cytotoxicity in a lung metastasis model. Second, B-cell maturation antigen (BCMA)-targeting DTX-adiposomes (BCMA@DTX-Ad) were formulated and showed improved therapeutic efficacy in a multiple myeloma mouse model. Finally, liver-targeting DTX-adiposomes (Liver@DTX-Ad) were developed and outperformed sorafenib in orthotopic liver cancer mouse models. Overall, our findings underpin the considerable potential of adiposome as a powerful drug delivery platform to treat various cancers. Notably, a series of adiposome targeting strategies was successfully established to achieve precise targeted delivery and better clinical efficacy, revealing that adiposome is a promising drug delivery platform with broad applicability.

## Results

### Construction and physicochemical characterization of DTX-Ad

To develop the DTX-Ad with high stability and encapsulation efficiency, DTX was first dissolved in selected neutral lipids (Fig. S1A-H) and then the mixture was transferred into an Eppendorf tube containing a phospholipid film followed by vigorous vortexing and centrifugation (Fig. 1A). The quality of blank adiposomes (without DTX) was determined by particle size, polydispersity index (PDI), and morphological observation (Fig. S1I-K). The physicochemical properties of DTX-Ad were comparable to those of blank adiposomes with an average diameter of 160 nm, PDI of 0.2, and zeta-potential value of -30 mV (Fig. 1B-C). DTX-Ad exhibited a uniform spherical morphology and no lipid aggregations and other contaminated structures were detected using fluorescence microscopy and transmission electron microscopy (TEM) (Fig. 1D-E). It was confirmed that DTX was encapsulated within the neutral lipid core of adiposomes by thin-layer chromatography (TLC) and high-performance liquid chromatography (HPLC), as DTX in adiposomes displayed identical migration distance in TLC and retention time in HPLC to the DTX standard (DTX-Std) and DTX-Inj (Fig. 1F-G; Fig. S1L). Further analysis found that the encapsulation efficiency of DTX-Ad was approximately 99.5% (Fig. 1H). The DTX-Ad formulation exhibited a uniformly milky appearance and remained physically stable for at least 321 days at 4°C, with minimal changes in particle size (150-170 nm) and DTX concentration (Fig. 1I-J). To examine the drug leakage under physiological conditions, DTX-Ad was dialyzed in phosphate-buffered saline (PBS) for 2 weeks under the condition of horizontal shaking. The average size of adiposomes was increased slightly to around 200 nm and the PDI remained unchanged, and the morphology remained as intact nanoparticles (Fig. 1K-M; Fig. S1M). In contrast, DTX in DTX-Inj was released rapidly during dialysis, whereas DTX-Ad maintained over 70% of its drug content after 2 weeks, indicating stable DTX retention (Fig. 1N). Collectively, DTX was efficiently encapsulated to adiposomes with long-term stability, providing a new water-soluble formulation of DTX for therapeutic use in cancer treatment.

**Figure 1.**
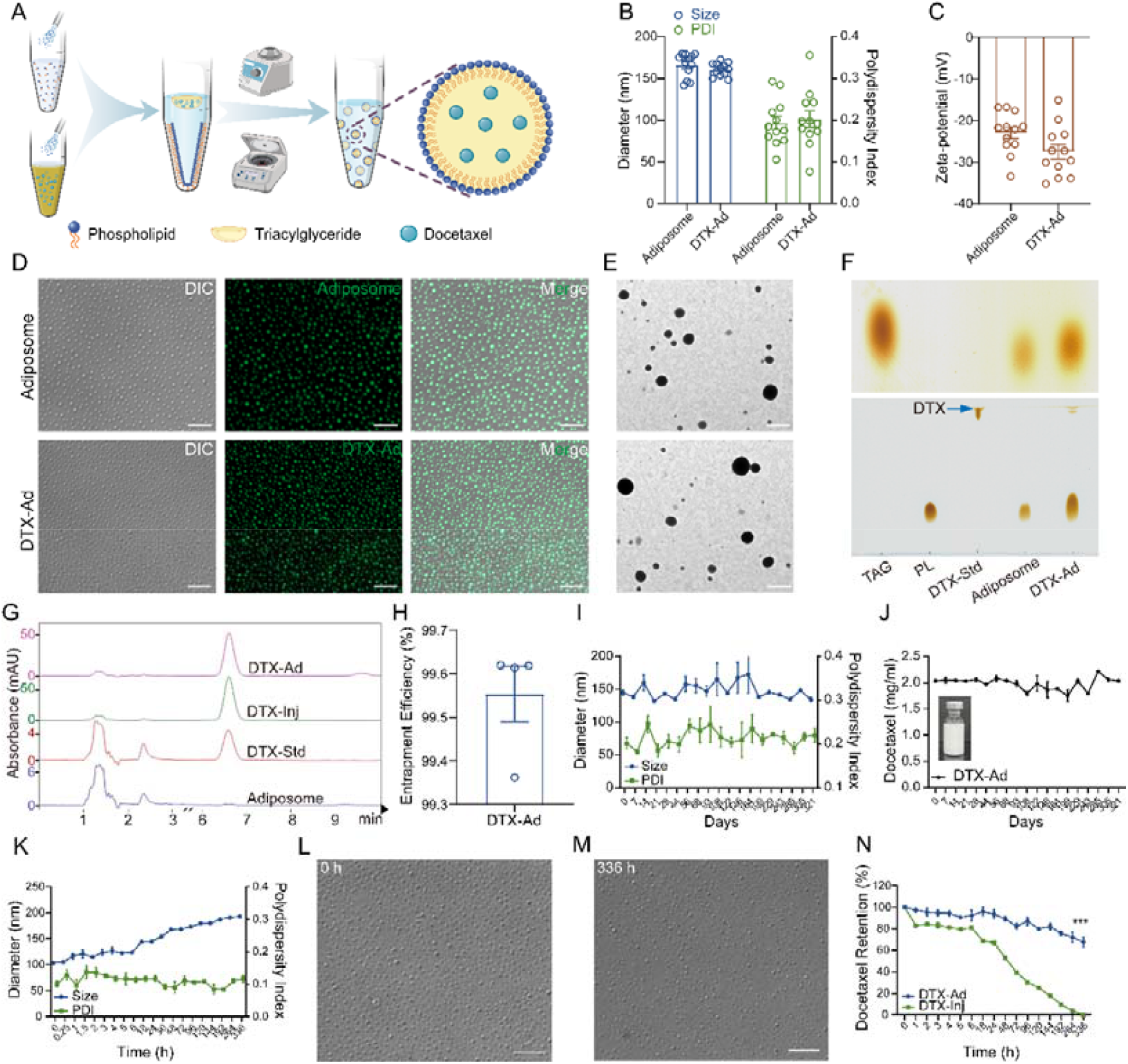
Construction and physicochemical characterization of DTX-Ad. **(A)** Schematic illustration of DTX-Ad preparation. DOPC in ethanol was added in an Eppendorf tube and ethanol evaporated under nitrogen gas. DOPC formed a phospholipid film on the surface of tube. In parallel, docetaxel (DTX) dissolved in ethanol was mixed with neutral lipids and the organic solvents evaporated under nitrogen gas to generate DTX-containing neutral lipids. Deionized water was added to hydrate the phospholipid film, followed by addition of the DTX-containing neutral lipids. After vortexing and centrifugation, DTX-Ad was obtained. **(B-C)** Particle size, PDI, and zeta-potential of blank adiposomes and DTX-Ad were measured by dynamic light scattering (DLS) (n = 12). **(D-E)** Images of adiposomes taken by fluorescence microscope and transmission electron microscope (TEM). For fluorescence imaging, adiposomes were mounted on glass slides and imaged directly. For TEM, adiposomes were positively stained before imaging. Scale bars: 5 μm (D) and 500 nm (E). **(F-G)** Thin-layer chromatography (TLC) and high-performance liquid chromatography (HPLC) analyses of DTX-Ad, compared with DTX standard (DTX-Std) and DTX-Inj. **(H)** Encapsulation efficiency of DTX in adiposomes was determined using HPLC to measure total and free DTX (n = 4). **(I-J)** Long-term stability of DTX-Ad during storage at 4°C was evaluated by DLS and HPLC (n = 3). Inset shows appearance of the DTX-Ad. **(K-M)** Stability of DTX-Ad during dialysis in PBS was determined by DLS and morphological observation at the indicated time points. Scale bar: 5 μm. **(N)** Retention of DTX-Ad and DTX-Inj after dialysis in PBS (n = 3). Data are presented as mean ± SEM. Statistical differences were calculated using two-tailed Student’s *t* test. \*\*\**P*<0.001.

### DTX-Ad inhibits microtubule depolymerization

Following physicochemical characterization, DTX-Ad was further evaluated for DTX-mediated microtubule stabilization and cytotoxicity in comparison with DTX-Inj. Consistent with DTX-Inj, DTX-Ad induced pronounced β-tubulin bundling with cytoskeletal collapse visualized by confocal microscopy (Fig. 2A). The form of β-tubulin displayed higher polymerization with stronger signal in the pellets after DTX-Ad treatment using centrifugation and Western blot analysis (Fig. 2B). Cell-cycle assay by flow cytometry showed that DTX-Ad-treated cells were predominantly arrested in G2 phase (Fig. 2C). Furthermore, DTX-Ad exhibited strong cytotoxicity among various tumor cells in a dose-dependent manner by CCK-8 assay, including PC-3, MCF-7, Hepa1-6 and H929, whereas blank adiposomes did not show any detectable cytotoxicity (Fig. 2D-G). Similar results were obtained in other cancer cell lines, including HEK293, MDA-MB-231, Huh-7, HeLa, and 4T1-Luc (Fig. S2A-E).

**Figure 2.**
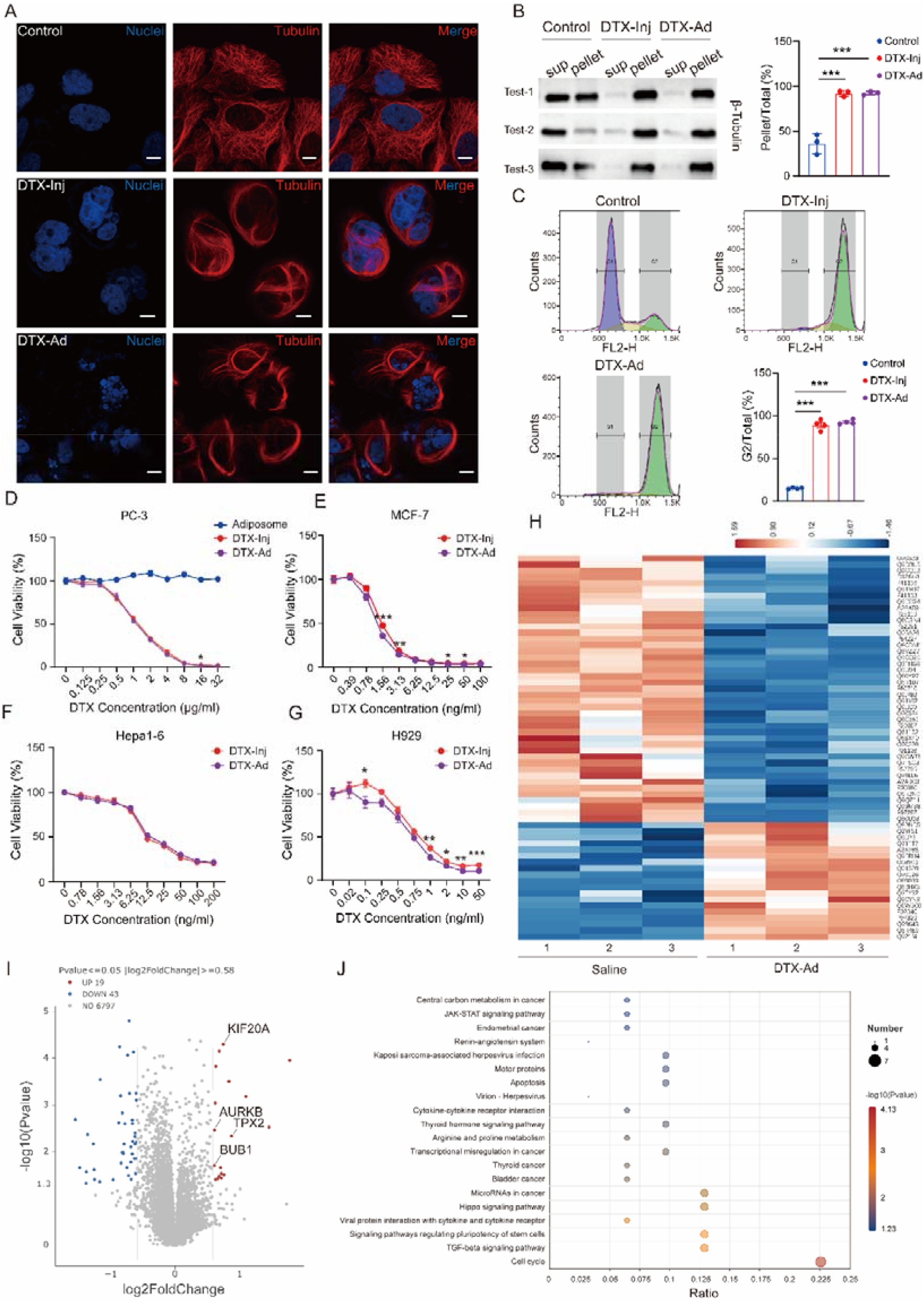
DTX-Ad induces microtubule stabilization and cell cycle arrest. **(A)** Confocal images of β-tubulin organization in MCF-7 cells treated with medium, DTX-Inj, or DTX-Ad at a DTX-equivalent concentration of 50 ng/mL for 24 h. Cells were stained with Hoechst for nuclei and Tubulin-Tracker Red for microtubules. Scale bars: 10 μm. **(B)** Western blot analysis of β-tubulin in supernatant (Sup) and pellet fractions from MCF-7 cells treated as in **(A)**, with quantification of the pellet-to-total ratio (n = 3). **(C)** Flow cytometry analysis of cell-cycle distribution in MCF-7 cells treated as in **(A)**, followed by ethanol fixation and propidium iodide staining (n = 4). **(D-G)** Cell viability of PC-3, MCF-7, Hepa1-6, and H929 cells treated with DTX-Inj, DTX-Ad, or blank adiposomes at the indicated concentrations, measured by CCK-8 assay. **(H)** Heatmap of differentially expressed proteins in Hepa1-6 cells treated with saline or DTX-Ad for 12 h at a DTX-equivalent concentration of 100 ng/mL (n = 3). **(I)** Volcano plot of differentially expressed proteins, with selected mitotic checkpoint-and spindle-associated proteins indicated. **(J)** KEGG pathway enrichment analysis of differentially expressed proteins. Data are presented as mean ± SEM. Statistical differences were calculated using two-tailed Student’s t test. \*\*\**P*<0.001, \*\**P*<0.01, \**P*<0.05.

To elucidate the cellular responses induced by DTX-Ad, proteomic analysis was performed. Heatmap analysis revealed substantial alterations in protein expression after DTX-Ad treatment (Fig. 2H). Volcano plot analysis identified significant upregulation of mitotic checkpoint and spindle-associated proteins, including BUB1, AURKB, TPX2, and KIF20A (Fig. 2I). KEGG enrichment analysis further showed that the most significantly affected pathways were associated with cell cycle regulation, apoptosis, and cancer-related signaling (Fig. 2J), supporting that DTX-Ad exerts its antitumor effects through canonical DTX-induced mitotic disruption. Additionally, DTX-Ad also modulated ER-associated protein processing and lipid-related pathways (Fig. S2F-H). Taken together, these results demonstrate that DTX-Ad exhibits robust cytotoxicity against various cancer cells by inhibiting microtubule depolymerization.

### DTX-Ad exhibits strong antitumor activity with less toxicity

To evaluate the efficacy and safety of DTX-Ad, three subcutaneous tumor models were established. In the H226 squamous cell carcinoma xenograft model, DTX-Ad and DTX-Inj significantly suppressed tumor growth with comparable efficacy (Fig. 3A). Notably, DTX-Ad treatment caused less weight loss than DTX-Inj (Fig. 3B), indicating that adiposome delivery improves systemic tolerance without compromising therapeutic potency. In the MCF-7 breast cancer xenograft model, DTX-Ad even displayed better antitumor efficacy than commercial albumin-bound paclitaxel (PTX-HSA) (Fig. 3C). More importantly, the subcutaneous tumors gradually disappeared with the treatment of DTX-Ad (Fig. 3D-F). DTX-Ad and PTX-HSA treatments exhibited comparable body weight gain, whereas DTX-Inj led to remarkable weight loss (Fig. 3G). DTX-Inj-treated mice showed significant decrease in blood neutrophil (NEU) count, while DTX-Ad treatment did not reduce NEU counts (Fig. 3H) or other blood cell populations, including white blood cells (WBC), lymphocytes (LYM), and monocytes (MON) (Fig. S3A-E). Additionally, similar results were observed following DTX-Ad treatment in the 4T1 syngeneic breast cancer model (Fig. S3F-J). Collectively, DTX-Ad outperforms PTX-HSA in tumor inhibition and exhibits less toxicity than DTX-Inj.

**Figure 3.**
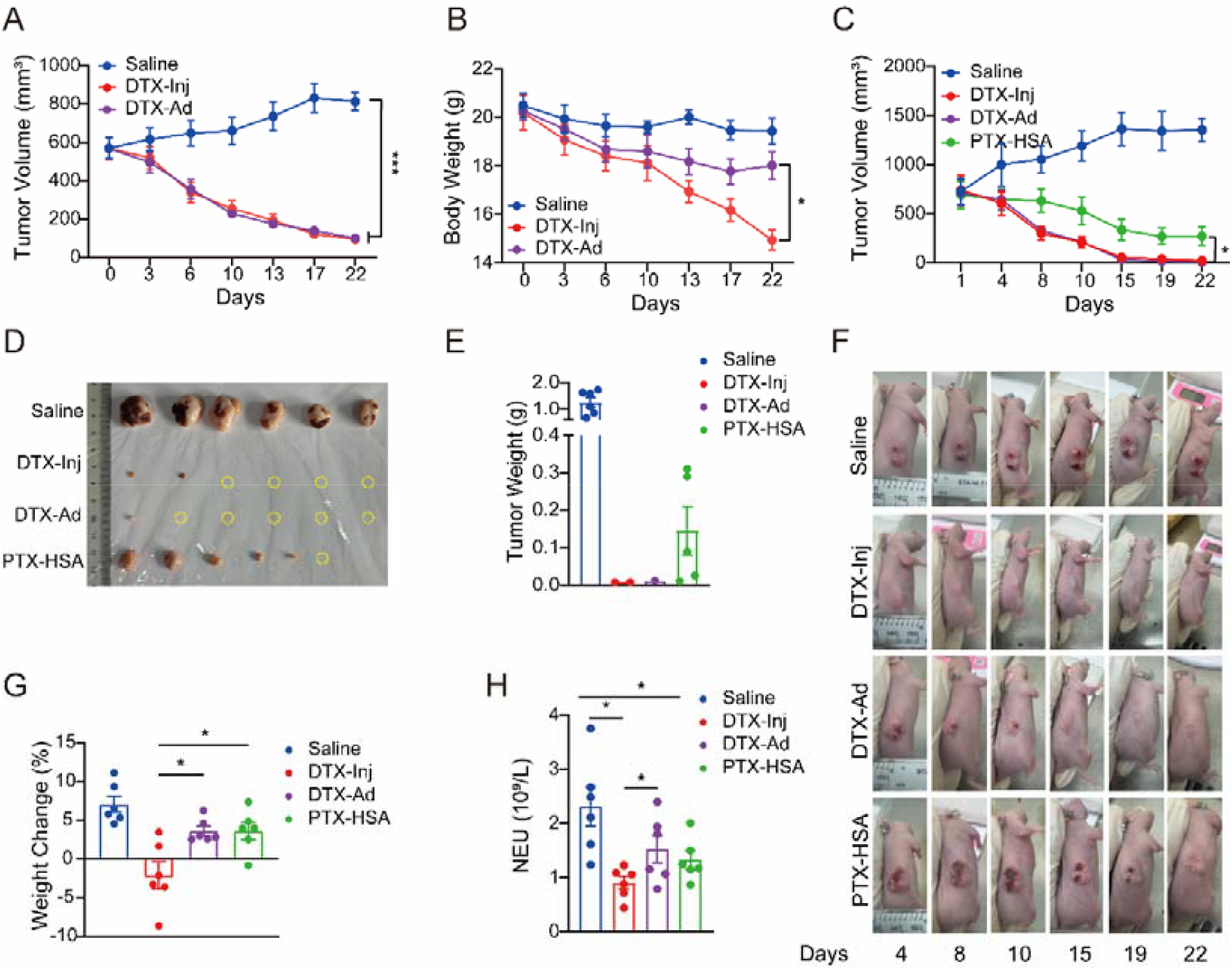
DTX-Ad exhibits better antitumor activity and tolerability than clinical formulations. **(A)** Tumor growth curves in the H226 xenograft model established by subcutaneous inoculation of 5 × 10^6^ H226 cells into BALB/c-nu mice, followed by treatment as indicated (n = 5). **(B)** Body weight change in H226 tumor-bearing mice during treatment. **(C)** Tumor growth curves in the MCF-7 xenograft model established by subcutaneous inoculation of 2 × 10^6^ MCF-7 cells into BALB/c-nu mice, followed by treatment as indicated (n = 6). **(D)** Representative images of excised tumors. **(E)** Weights of excised tumors. **(F)** Representative photographs of tumor-bearing mice during treatment. **(G)** Body weight change at the endpoint. **(H)** Blood neutrophil (NEU) counts in different groups. Data are presented as mean ± SEM. Statistical differences were calculated using two-tailed Student’s t test. \*\*\**P*<0.001, \**P*<0.05.

### Pep33 modification provides lung-targeting capability to adiposomes and enhances antitumor efficacy

Although DTX-Ad exhibited improved safety, its antitumor efficacy was comparable to DTX-Inj. To further enhance therapeutic performance, we designed a tissue-targeting adiposome formulation. Peptides containing the C-terminal R/KXXR/K motif have been reported to bind neuropilin-1 (NRP1) and enhance tumor penetration of nanocarriers^[33-35]^. AAM-B specifically localizes to the periphery of lipid droplets via hydrophobic targeting sequences^[36]^. Thus, Lung@DTX-Ad was developed by fusing AAM-B with the CRGDK motif, forming a 33-amino-acid peptide (Pep33) (Fig. 4A). Pep33 attaches to the surface of adiposomes via AAM-B and enables lung targeting through CRGDK binding to NRP1. Confocal imaging showed that Pep33-EGFP formed a ring structure surrounding lipid droplets, whereas EGFP was diffusely distributed in the cytosol, indicating that Pep33 specifically recognizes the phospholipid monolayer of lipid droplets (Fig. 4B). We next tested whether Pep33 attaches to adiposomes after incubation and purification. Silver staining confirmed successful binding of Pep33 to the adiposomes (Fig. 4C). Although Pep33 loading led to modest decreases in particle size, PDI, and zeta-potential magnitude, the overall physicochemical stability of adiposomes was preserved (Fig. 4D-E).

**Figure 4.**
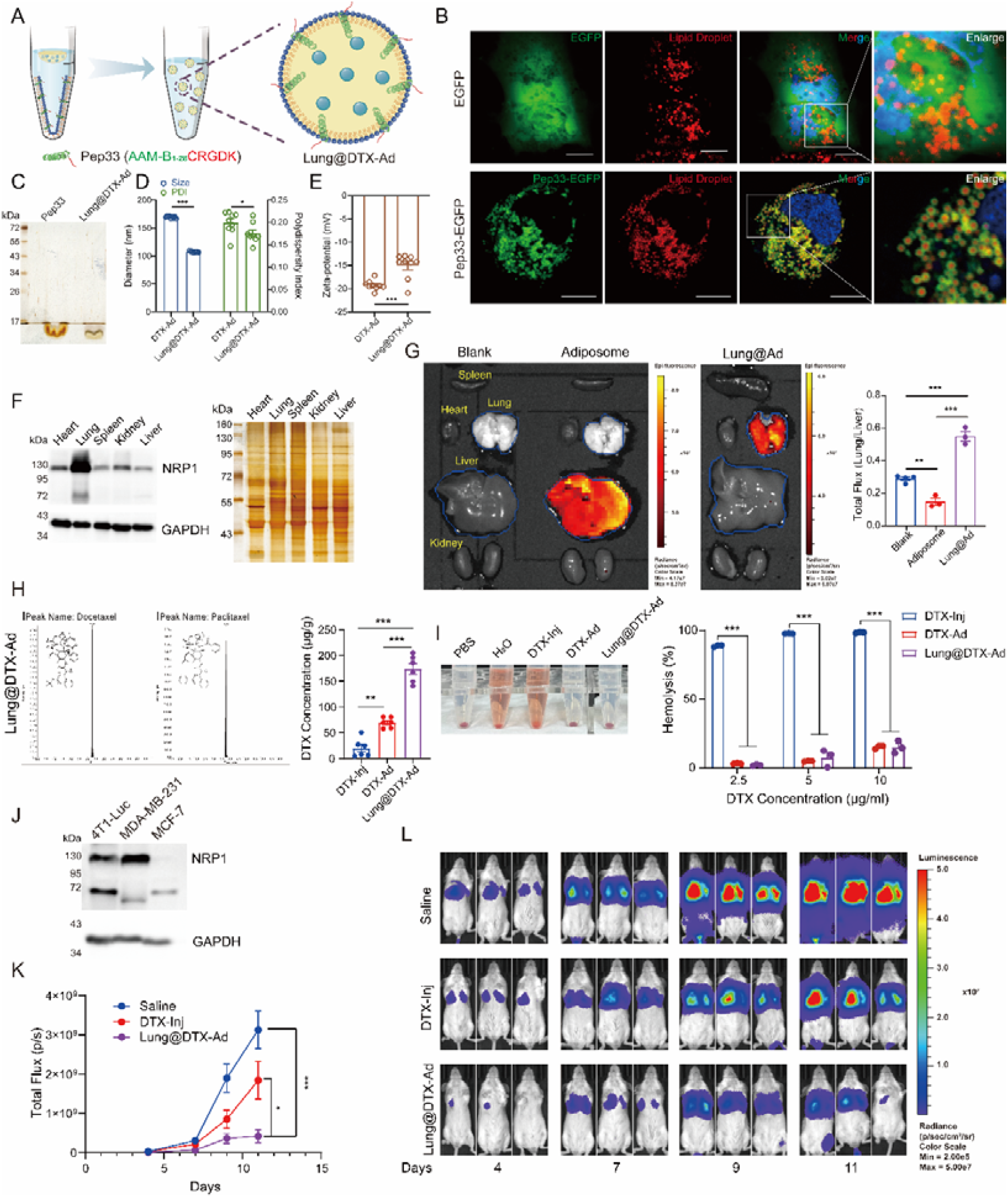
Pep33 functionalization confers lung-targeting capability to adiposomes and enhances therapeutic efficacy. **(A)** Schematic illustration of Lung@DTX-Ad. Pep33 contains an NRP1-binding motif (CRGDK, red) and a lipid droplet-targeting sequence (AAM-B, green), and was incorporated during adiposome preparation by co-evaporation with DOPC. **(B)** Confocal images of cells transfecting EGFP or Pep33-EGFP. Lipid droplets were stained with LipidTOX Red. Scale bars: 10 μm. **(C)** Silver staining analysis of Pep33 associated with adiposomes after purification. **(D-E)** Physicochemical characterization of DTX-Ad with or without Pep33 modification, measured by DLS (n = 9). **(D)** Western blot and silver staining analyses of NRP1 expression in major organs. **(E)** *Ex vivo* fluorescence imaging of major organs collected 3 h after intravenous injection of saline, fluorescent adiposomes, or Lung@Ad into BALB/c mice, with quantification of the lung-to-liver signal ratio (n = 3 or 4). **(F)** DTX concentrations in lung tissues measured by LC-MS/MS at 1 h after intravenous injection of the indicated formulations (n = 6). **(H)** Hemolysis assay of different formulations at the indicated concentrations. **(I)** Western blot analysis of NRP1 expression in different cancer cell lines. **(J)** Quantification of tumor burden in a lung metastasis model established by intravenous injection of 5 × 10^4^ 4T1-Luc cells into BALB/c mice, followed by treatment as indicated (n = 7). **(K)** Representative bioluminescence images of mice during treatment. Data are presented as mean ± SEM. Statistical differences were calculated using two-tailed Student’s t test. \*\*\**P*<0.001, \*\**P*<0.01, \**P*<0.05.

Since NRP1 was most abundantly expressed in the lung among major organs (Fig. 4F), we further evaluated the *in vivo* distribution of Pep33-modified adiposomes (Lung@Ad). Compared to blank adiposomes enriched in the liver, Lung@Ad selectively accumulated in the lung (Fig. 4G). Consistent with this observation, the DTX concentration in the lung delivered by Lung@DTX-Ad was 8.7-fold higher than that delivered by DTX-Inj at an equivalent DTX dose (Fig. 4H). In addition, Lung@DTX-Ad barely caused hemolysis compared to severe hemolysis induced by DTX-Inj (Fig. 4I). To evaluate the therapeutic efficiency of Lung@DTX-Ad *in vivo*, 4T1-Luc with high expression of NRP1 was chosen to establish a lung-metastasis model (Fig. 4J). Lung@DTX-Ad exhibited the strongest suppression of pulmonary tumor burden (Fig. 4K-L). These results showed that Lung@DTX-Ad selectively delivered DTX to lung tumor-bearing tissues and achieved improved therapeutic efficacy.

### BCMA antibody modification confers multiple myeloma-targeting capability to adiposomes and achieves sustained suppression

Antibody-drug conjugates (ADCs) represent a novel class of biopharmaceuticals combining a tumor-specific monoclonal antibody with potent cytotoxic agents via an engineered linker. Despite their therapeutic benefits, ADCs face significant challenges, including linker instability and insufficient payload^[37;38]^. To address the aforementioned problems, antibody-conjugated nanoparticles (ACNPs) have been developed^[39]^. Here, BCMA@DTX-Ad was constructed using the biotin-avidin system (Fig. 5A). Biotin-labeled adiposomes (Bio-Ad) were initially constructed by adding biotin-conjugated phospholipid (Biotin-DOPE) into the phospholipid mixture during preparation (Fig. S4A-D). GFP-Avidin fusion protein (tagged at the C-terminal end of GFP, Fig. S4E) formed a ring-like signal on Bio-Ad but not on unmodified adiposomes, confirming the feasibility of the system (Fig. 5B). The BCMA-ab-Avidin (tagged at the heavy chain of BCMA antibody) was purified (Fig. S4F) and incubated with Bio-Ad. BCMA-ab-Avidin was efficiently recruited onto the Bio-Ad in a biotin dose-dependent manner (Fig. 5C). Despite modest variations in particle size and zeta-potential following surface functionalization, the adiposomes maintained their structural integrity and colloidal stability (Fig. 5D-E).

**Figure 5.**
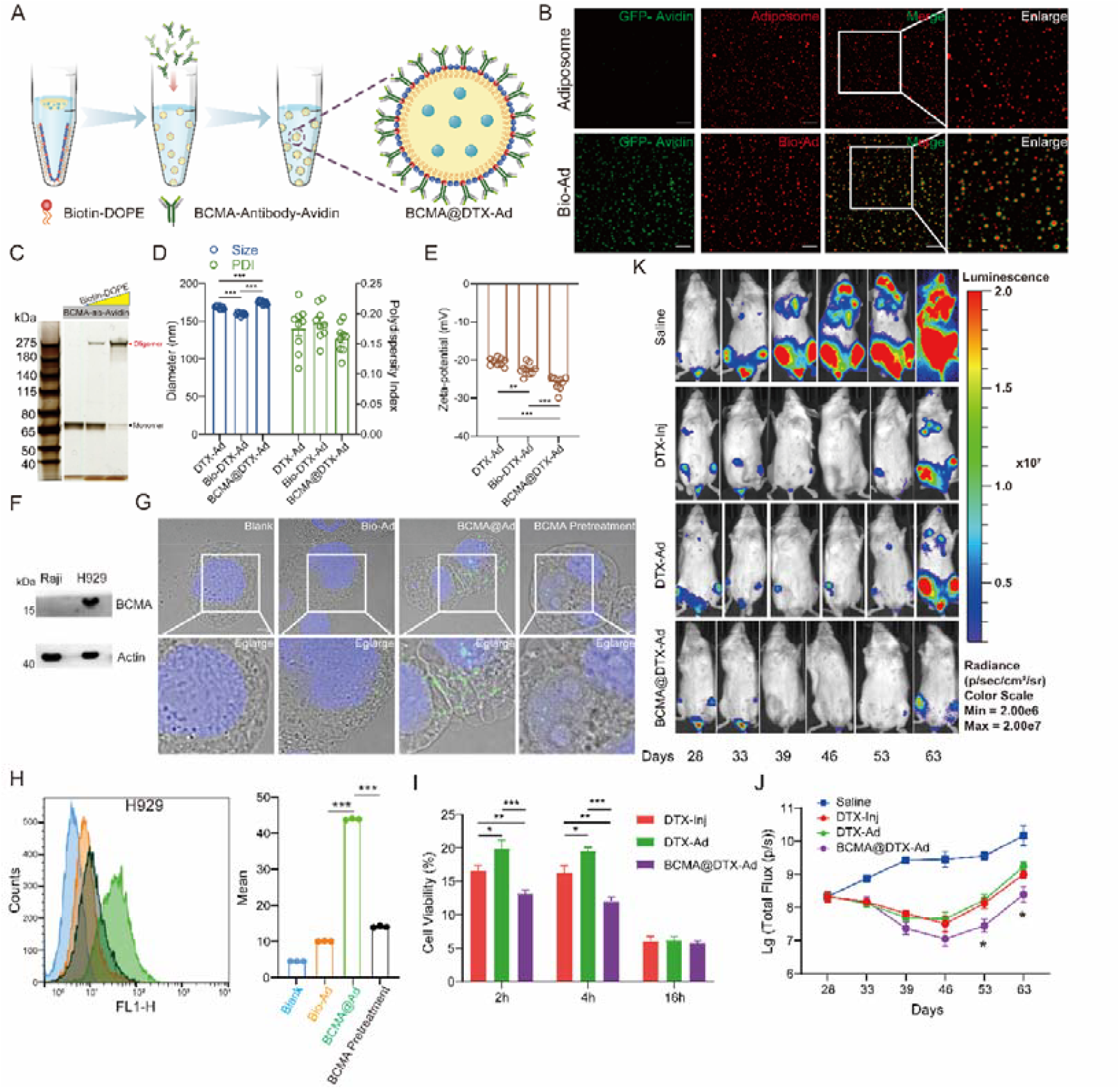
BCMA antibody modification enables multiple myeloma targeting and sustained antitumor efficacy. **(A)** Schematic illustration of BCMA@DTX-Ad constructed through biotin-avidin interaction. **(B)** Confocal images of GFP-avidin localization on Bio-Ad and unmodified adiposomes. Scale bars: 5 μm. **(C)** Silver staining analysis of BCMA-ab-Avidin associated with adiposomes in the presence or absence of biotin-DOPE. **(D-E)** Physicochemical characterization of adiposomes after biotin-DOPE incorporation and BCMA antibody functionalization, measured by DLS (n = 9). **(F)** Western blot analysis of BCMA expression in multiple myeloma cell lines. **(G)** Confocal images of BCMA@Ad uptake in H929 cells with or without preincubation with recombinant BCMA protein. **(H)** Flow cytometry analysis of BCMA@Ad uptake in H929 cells treated with the indicated formulations (n = 3). **(I)** Cell viability of H929 cells treated with the indicated formulations for the indicated durations. **(J-K)** *In vivo* bioluminescence imaging and quantification of tumor progression in a systemic multiple myeloma model established by intravenous injection of 1 × 10^7^ U266-Luc cells into NCG mice, followed by treatment as indicated (n = 8). Data are presented as mean ± SEM. Statistical differences were calculated using two-tailed Student’s *t* test. \*\*\**P*<0.001, \*\**P*<0.01, \**P*<0.05.

Then, the H929 multiple myeloma cells with high BCMA expression (Fig. 5F) were chosen to test the targeting of BCMA antibody-bound adiposomes (BCMA@Ad). Compared to adiposomes without BCMA antibody, BCMA@Ad showed greater internalization by H929 cells. Meanwhile, Preincubation of BCMA@Ad with recombinant BCMA protein reduced its binding/uptake by H929 cells (Fig. 5G). Flow cytometry analysis also demonstrated greater uptake of BCMA@Ad by H929 cells than Bio-Ad and BCMA-pretreated adiposomes (Fig. 5H). In Raji cells without BCMA expression, BCMA@Ad displayed comparable signal to Bio-Ad (Fig. S4G). In addition, BCMA@DTX-Ad exhibited higher cytotoxicity against H929 cells after treatment for 2 h and 4 h (Fig. 5I). Moreover, in the U266-Luc multiple myeloma mouse model with high BCMA expression (Fig. S4H), BCMA@DTX-Ad inhibited tumor growth and achieved sustained suppression after discontinuing the treatment (Fig. 5J-K). These results indicate the feasibility of antibody-modified adiposomes.

### ApoE modification presents liver-targeting capability to adiposomes and augments therapeutic effect

Intravenously administered nanoparticles are mainly accumulated in the liver due to the passive accumulation and uptake by liver macrophages^[40]^. Active delivery of nanoparticles to specific liver cell types remains challenging. Based on our previous research using adiposomes to construct liver-targeting artificial lipoproteins^[25]^, Liver@DTX-Ad was developed (Fig. 6A) by attaching apolipoprotein E (ApoE), which is the ligand of low-density lipoprotein receptor (LDLR). The ApoE on the adiposomes was detected by the ring structure on the surface of adiposomes and protein gel electrophoresis (Fig. 6B-C). ApoE functionalization increased the absolute zeta-potential, indicating maintained colloidal stability (Fig. 6D-E).

**Figure 6.**
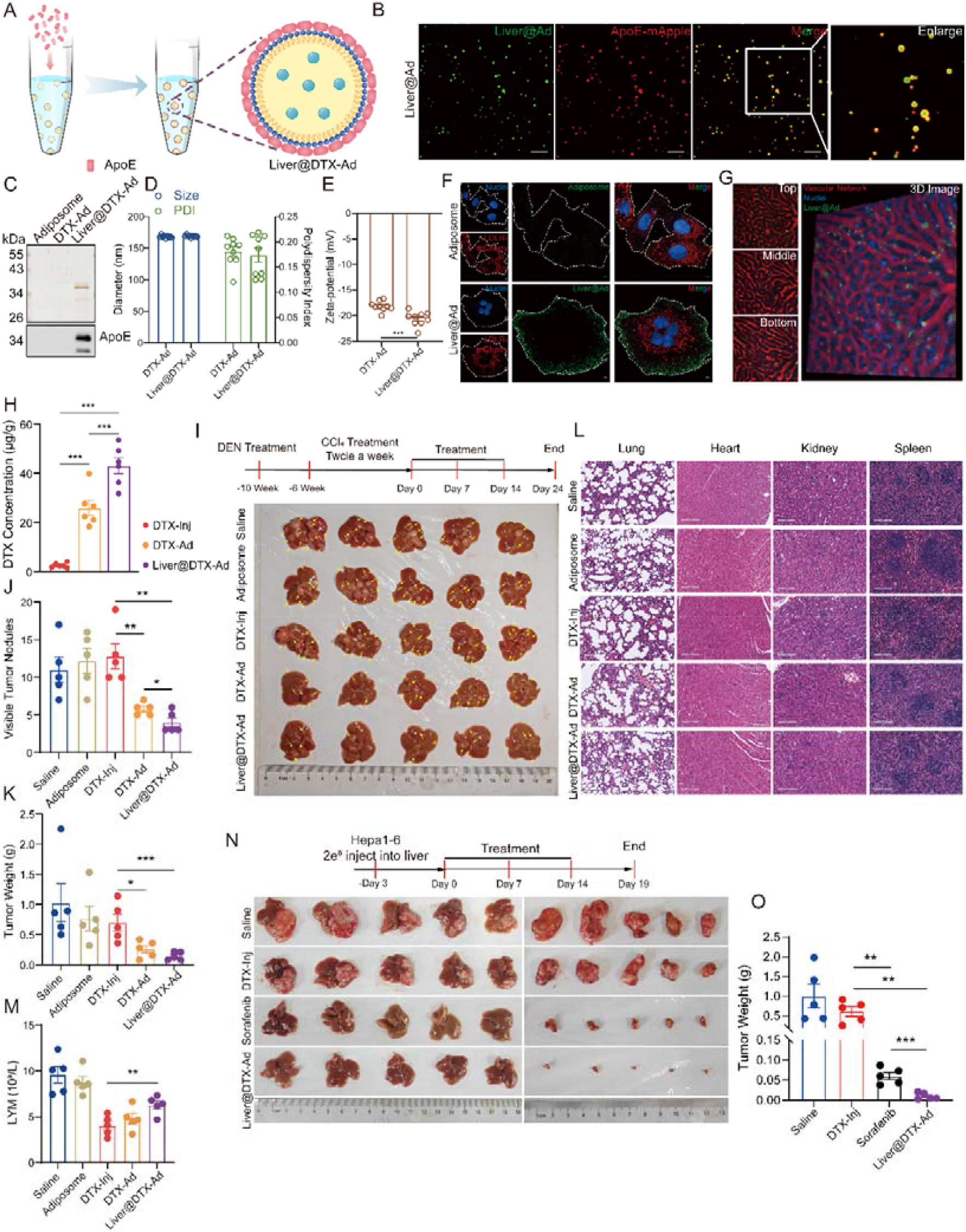
ApoE functionalization enables LDLR-mediated liver targeting and enhances therapeutic efficacy in hepatocellular carcinoma models. **(A)** Schematic illustration of Liver@DTX-Ad for LDLR-mediated liver targeting. ApoE was incubated with preformed adiposomes to generate Liver@DTX-Ad. **(B)** Confocal images of ApoE localization on adiposomes. Scale bars: 5 μm. **(C)** Silver staining and Western blot analysis of ApoE associated with DTX-Ad. **(D-E)** Physicochemical characterization of adiposomes with or without ApoE modification, measured by DLS (n = 9). **(F)** Confocal images of cellular uptake of Liver@Ad in LDLR-expressing HEK293 cells after incubation for 1 h under serum-free conditions. Scale bars: 10 μm. **(G)** Three-dimensional *in vivo* fluorescence imaging and sectional views after administration of Liver@Ad. **(H)** Docetaxel concentrations in liver tissues measured by LC-MS/MS at 1 h after intravenous injection of the indicated formulations (n = 6). **(I)** Experimental timeline and representative images from the DEN/CCl_4_-induced hepatocarcinoma model (n = 5). **(J-K)** Quantification of tumor burden in the hepatocarcinoma model, including visible tumor nodules and tumor weight. **(L)** Histological analysis of major organs in the indicated groups. **(M)** Hematological analysis of lymphocyte counts in the indicated groups. **(N)** Representative images from the orthotopic Hepa1-6 liver tumor model established by intrahepatic injection of 2 × 10^6^ Hepa1-6 cells mixed with Matrigel, followed by treatment as indicated (n = 5). **(O)** Tumor weights in the orthotopic Hepa1-6 model. Data are presented as mean ± SEM. Statistical differences were calculated using two-tailed Student’s *t* test. \*\*\**P*<0.001, \*\**P*<0.01, \**P*<0.05.

Then, the targeting of ApoE-coated adiposomes was evaluated *in vitro* and *in vivo*. ApoE-coated adiposomes (Liver@Ad) were taken up more in LDLR-overexpressing cells than blank adiposomes (Fig. 6F). Real-time *in vivo* three-dimensional fluorescence imaging revealed preferential accumulation of Liver@Ad in the liver (Fig. 6G, Video. 1-2). Consistent with this observation, the DTX concentration in the liver delivered by Liver@DTX-Ad was 16.8-fold higher than that delivered by DTX-Inj (Fig. 6H). Next, the spontaneous hepatocarcinoma mouse model induced by diethylnitrosamine (DEN) and carbon tetrachloride (CCl_4_) was constructed to evaluate the therapeutic efficacy of Liver@DTX-Ad. The Liver@DTX-Ad exhibited the best antitumor activity with the fewest tumor nodules and the lowest tumor weight (Fig. 6I-K). Histological analysis showed no obvious tissue damage in the Liver@DTX-Ad groups (Fig. 6L), indicating good systemic tolerability. Serum biochemical analysis showed that total cholesterol, HDL-C, and LDL-C levels remained normal after Liver@DTX-Ad administration (Fig. S5A-C). In addition, Liver@DTX-Ad preserved lymphocyte counts, whereas DTX-Inj reduced WBC and NEU levels (Fig. 6M; Fig. S5D-E), suggesting improved hematological safety. Furthermore, the Hepa1-6 hepatoma cell line with high expression of LDLR (Fig. S5F) was orthotopically implanted in the liver to test the efficacy of Liver@DTX-Ad in comparison with the clinically used drug sorafenib. As a result, Liver@DTX-Ad significantly inhibited tumor growth and even achieved tumor eradication (Fig. 6N), outperforming DTX-Inj and sorafenib (Fig. 6O). These results indicate that Liver@DTX-Ad achieves active targeting to the liver with enhanced efficacy and reduced systemic toxicity.

## Discussion

The delivery of hydrophobic small-molecule drugs remains a central challenge in therapeutic areas and DTX exemplifies this dilemma. Despite the tremendous efforts to develop various delivery methods to address the limitations of DTX over the past several decades^[41;42]^, Taxotere® and its generic solvent-based injections remain the only clinical formulations of DTX^[28]^. Taxotere® is associated with serious side effects, including acute hypersensitivity reactions, cumulative fluid retention and asthenia due to the presence of Tween 80 and ethanol^[32; 43]^. With the emergence of nanotechnology, various nanoparticles have been utilized to deliver DTX owing to their tunable size, surface modifiability, and enhanced permeability and retention (EPR) effect^[44]^. Nevertheless, these formulations still face challenges in biocompatibility, pharmacokinetics, and tumor-specific targeting. Inspired by the natural lipid-transport machinery of lipid droplets and lipoproteins, adiposome was engineered to improve the clinical benefits of DTX.

DTX-Ad without surface modification was initially generated and exhibited improved safety, as evidenced by reduced body-weight loss, and preserved neutrophil counts (Fig. 3G-H). The favorable safety of DTX-Ad is attributable to its naturally derived lipid composition, such as phospholipids and triacylglycerols^[45]^. In addition, the chosen triacylglycerols consist of a glycerol backbone esterified with unsaturated fatty acids that have been widely considered beneficial^[46; 47]^. Beyond oncology, adiposome could be precisely designed to encapsulate functional lipids for the treatment of metabolic diseases, such as omega-3 fatty acids, sphingolipids, and retinoic acid^[48-50]^.

Given that non-targeted DTX-Ad displayed comparable antitumor activity to DTX-Inj, we creatively designed targeted adiposomes using different approaches to achieve desirable therapeutic effects. Based on proteomic analyses of purified lipid droplets from diverse organisms^[51; 52]^, certain LD-resident proteins specifically localize to the lipid droplet monolayer membrane^[21; 53; 54]^, such as AAM-B and PLIN2 in mammals, MDT-28 and DHS-3 in *C. elegans*, and microorganism lipid droplet small (MLDS) in bacteria. Owing to a shared but incompletely understood recognition for the phospholipid monolayer, these proteins can be harnessed to anchor ligands of interest onto the adiposome surface. In our work, Lung@Ad were developed by attaching a 33-amino-acid fusion peptide (Pep33) comprising AAM-B_1-28_, which anchors the adiposome surface, and the CendR-related CRGDK motif capable of engaging NRP1 highly expressed in the lung^[55]^ (Fig. 4F). The feasibility of Lung@Ad was demonstrated through *in vivo* imaging and drug tissue distribution (Fig. 4G-H). Importantly, Lung@DTX-Ad exhibited enhanced antitumor activity in a lung metastasis model (Fig. 4L). Given the growing global burden of respiratory diseases driven by air pollution^[56]^, Lung@Ad represents a potent delivery platform for diverse pulmonary conditions, including asthma, chronic obstructive pulmonary disease (COPD), and acute respiratory infections.

Beyond peptide-based targeting, antibodies were further employed to confer high specificity to adiposome while overcoming the limitations of conventional ADCs. Antibodies are highly specific targeting ligands that have been widely employed in the research and clinical application of ADCs^[57]^. However, conventional ADCs are subject to inherent limitations, including a low drug-to-antibody ratio and unstable linker cleavage resulting in off-target toxicity^[58; 59]^. To overcome these shortcomings, ACNPs have emerged as a highly promising alternative strategy^[60]^. Here, antibody-conjugated adiposome was developed through a biotin-avidin system (Fig. 5A). The resulting BCMA@DTX-Ad enhanced BCMA-dependent uptake and prolonged tumor suppression in a multiple myeloma mouse model (Fig. 5K). Compared with conventional ADCs, the antibody-functionalized adiposome offers distinct advantages, including enhanced binding stability with reduced off-target risk, superior drug-loading capacity, and synergistic effect of multiple antibodies. Collectively, the antibody-functionalized adiposome delivery system demonstrates broad prospects for clinical application.

The liver-targeting strategy leverages the endogenous lipoprotein-receptor axis that originally inspired the adiposome design. The liver is the largest metabolic organ in the body^[61]^, comprising various cell types that perform distinct functions, including hepatocytes (approximately 60%), hepatic stellate cells, liver sinusoidal endothelial cells, and Kupffer cells. Intravenously administered nanocarriers tend to passively accumulate in the liver, primarily owing to the dramatically reduced blood flow velocity within hepatic sinusoids ^[62]^ and substantial engulfment of Kupffer cells^[63]^. Therefore, active targeting to specific hepatic cell subpopulations is of critical importance for the precise treatment of various liver diseases^[64; 65]^. Our previous work further demonstrated that apolipoprotein-functionalized adiposomes can be used to construct artificial lipoproteins with biological activity^[66]^. In the present study, Liver@DTX-Ad was developed by surface decoration with ApoE, a key apolipoprotein ligand for LDLR-family receptors^[67; 68]^ (Fig. 6A). The Liver@DTX-Ad demonstrated significantly superior antitumor efficacy compared with DTX-Inj and the clinically used multi-kinase inhibitor sorafenib in multiple hepatocellular carcinoma mouse models (Fig. 6I, N). Therefore, the Liver@Ad platform may be further extended to the diagnosis and treatment of chronic liver diseases.

Taken together, the present study provides a biomimetic strategy of delivering hydrophobic drugs in an aqueous physiological environment. The nature-inspired adiposome efficiently encapsulates DTX while eliminating potentially toxic excipients present in the commercial formulation (Tween 80 and ethanol), and achieves precise tumor targeting with improved therapeutic outcomes across lung cancer, multiple myeloma, and hepatocellular carcinoma models. Our team is actively accelerating the clinical translation of the DTX-loaded adiposome formulation. Beyond small-molecule drugs, adiposome serves as a transformative delivery platform with the potential to advance the therapeutic application of biological drugs.

## Acknowledgments

The authors thank Dr. Yanan Gao for her help in the establishment of the tumor-bearing mouse model. The authors thank Ms. Yan Teng and Ms. Chunliu Liu (Center for Biological Imaging, IBP, CAS) for their help in taking and analyzing confocal images. The authors thank Mr. Junying Jia and Ms. Shu Meng (Core Facility, Institute of Biophysics, Chinese Academy of Sciences) for the technical support in flow cytometry analyses. The authors thank Ms. Weihua Wang in the Center of Pharmaceutical Technology. Tsinghua University, for LC-MS/MS analysis. The authors thank Yihui Xu (State Key Laboratory of Biomacromolecules, Institute of Biophysics, Chinese Academy of Sciences) for in vivo imaging technology. The authors thank Nanjing Super Vision Imaging Technology Co., Ltd. for providing the JP-01, and Walker Luo for technical assistance in image acquisition. This work was supported by the National Key Research and Development Program of China (Grant No. 2023YFA1801103 and 2024YFA1306101), National Natural Science Foundation of China (Grant No. 92357302), Beijing Research Ward Excellence Program (Grant No. BRWEP2024W102170106), Beijing Natural Science Foundation (Grant No. L252192 and 7244522), and the Prevention and Control of Emerging and Major Infectious Diseases-National Science and Technology Major Project (Grant No. 2025ZD01906004).

## Author Contributions

B.P., Z.C., and P.L. conceived and designed the study. B.P. and Z.C. performed most of the experiments, analyzed the data, prepared the figures, and wrote the original draft. Z.L., G.Z., C.Z., Z.Z., Y.Q.Z., Q.Z., S.L., Y.H.Z, B.Y., X.L., and K.X.L. contributed to adiposome preparation, physicochemical characterization, cell experiments, animal experiments, imaging analysis, or data collection. S.Z. assisted with electron microscopy analysis. P.L. and Z.C. supervised the project, provided resources and funding support, and revised the manuscript. All authors discussed the results and approved the final manuscript.

## Materials and Methods

### Materials

Docetaxel powder was purchased from Wuxi Taxus Pharmaceutical Co., LTD. Commercial docetaxel injection was purchased from Yangtze River Pharmaceutical Group. Commercial albumin-bound paclitaxel was purchased from CSPC Pharmaceutical Group. Pep33 was custom made from GL Biochem (Shanghai) Ltd with the amino acid sequence MELTIFILRLAIYILTFPLYLLNFLGLWCRGDK. BALB/c and C57BL/6J mice were purchased from SPF Biotechnology Co., Ltd. BALB/c-nu and NCG mice were obtained from GemPharmatech Co., Ltd. All mice were maintained under specific pathogen-free (SPF) conditions with a 12 h light-dark cycle and free access to food and water. All animal experiments were approved by the Laboratory Animal Management and Use Committee of the Institute of Biophysics, Chinese Academy of Sciences (Approval No. SYXK2023169). U266-luc cells were kindly provided by Prof. Shengdian Wang, and Hepa1-6 cells were kindly provided by Prof. Pengyuan Yang. Other cells were obtained from the American Type Culture Collection (ATCC). All cells were cultured under standard conditions according to recommended protocols.

### Preparation of adiposome formulations

Adiposomes were prepared according to a previously established protocol with minor modifications^[25]^. Briefly, 2 mg 1,2-dioleoyl-sn-glycero-3-phosphocholine (DOPC, Avanti Polar Lipids, #850375) dissolved in ethanol was dried under nitrogen to form a phospholipid film. 5 μL neutral lipids consisting of fish oil (Aladdin, #F1456055) and tricaprylin (Aladdin, #T1521612) (1:1, v/v) were then added and blank adiposomes were generated by vortex-assisted self-assembly in the aqueous phase, followed by sequential centrifugation purification.

For preparation of DTX-Ad, docetaxel (20 mg/mL in ethanol) was mixed with neutral lipids at an equal volume ratio, followed by nitrogen evaporation to remove ethanol, yielding docetaxel-containing neutral lipids. All subsequent steps were identical to those used for blank adiposomes.

For preparation of Lung@Ad or Lung@DTX-Ad, Pep33 (10 μg) was dissolved in the same solvent mixture as DOPC and co-evaporated prior to adiposome assembly.

For preparation of BCMA@Ad or BCMA@DTX-Ad, Biotin-labeled adiposomes (Bio-Ad) were generated by incorporating 1,2-dioleoyl-sn-glycero-3-phosphoethanolamine-N-(cap biotinyl) (Biotin-DOPE, Avanti Polar Lipids, #870273) and DOPC at a mass ratio of 1:7 during phospholipid film formation. The resulting Bio-Ad was incubated with avidin-fusion proteins (GFP-avidin or BCMA antibody-avidin) at a final protein concentration of 0.08 mg/mL for 1 h at room temperature with gentle mixing every 10 min. Unbound proteins were removed by centrifugation and washing.

For preparation of Liver@Ad or Liver@DTX-Ad, purified ApoE or ApoE-mApple was incubated with adiposomes at a final concentration of 0.15 mg/mL for 1 h at room temperature, followed by washing to remove free protein.

### Morphological observation

For morphological observation, adiposome suspensions were diluted with ultrapure water and mounted onto glass slides, followed by gentle placement of coverslips. Unlabeled adiposomes were imaged using an optical microscope (ZEISS Axio Imager M2). Fluorescent adiposomes were imaged using the same microscope under fluorescence mode or a confocal laser scanning microscope (OLYMPUS FV3000RS). Representative images were collected for morphological analysis. For TEM, purified adiposome suspensions were deposited onto copper grids and allowed to adsorb. The grids were fixed with 2.5% glutaraldehyde (0.1 M phosphate buffer, pH 7.2) for 10 min, followed by staining with 1% osmium tetroxide, 0.1% tannic acid, and 2% uranyl acetate for 10 min each, with water washes between staining steps. After air drying, samples were imaged using a CM120-FEG transmission electron microscope (FEI) operated at 120 kV.

### High-performance liquid chromatography

Docetaxel and paclitaxel contents were quantified by HPLC. For sample preparation,

10 μL of DTX-Ad, DTX-Inj, or PTX-HSA was mixed with 190 μL methanol, vortexed thoroughly, and centrifuged at 21,380 × *g* for 3 min. The supernatant was collected, and 10 μL was injected for analysis. Chromatographic separation was performed using a ZORBAX StableBond column (Agilent, #883975-902). The mobile phase consisted of methanol/acetonitrile/water (4:3:3, v/v/v), delivered at a flow rate of 1.0 mL/min. The column temperature was maintained at 30°C, and analytes were detected at 227 nm. Standard curves were generated using docetaxel standards under identical conditions, and drug concentrations in samples were calculated according to the corresponding peak areas.

### Encapsulation efficiency determination

Total docetaxel content in DTX-Ad was first measured directly. To determine free docetaxel, 250 μL of DTX-Ad was transferred into a centrifugal concentrator tube (Nanosep, #OD300C35) and centrifuged at 10,000 × g for 10 min. The filtrate volume was recorded, and free docetaxel concentration was measured by HPLC. Encapsulation efficiency (EE) was calculated as:

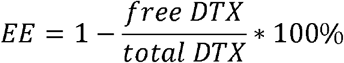

### Microtubule polymerization and cell cycle analysis

MCF-7 cells were seeded into six-well plates and cultured overnight until approximately 70% confluence. Cells were treated with DTX-Inj or DTX-Ad at a docetaxel-equivalent concentration of 50 ng/mL for 24 h. For microtubule polymerization analysis, cells were preincubated at 4°C and collected by trypsinization. Cells were washed with prechilled PBS and incubated with 100 μL hypotonic buffer (1 mM MgCl_2_, 2 mM EGTA, 0.5% Nonidet P-40, 2 mM phenylmethylsulfonyl fluoride, 200 units/mL aprotinin, 100 μg/mL soybean trypsin inhibitor, 5.0 mM ε-amino caproic acid, 1 mM benzamidine, and 20 mM Tris-HCl, pH 6.8) at 37°C for 5 min to allow separation of soluble and polymerized tubulin fractions. Samples were centrifuged at 21,380*g* for 10 min at 4°C. Supernatants containing soluble tubulin were collected and mixed with an equal volume of 2× sample buffer (125 mM Tris-HCl, 20% glycerol, 4% SDS, 4% β-mercaptoethanol, 0.04% bromophenol blue, pH 6.8). Pellets containing polymerized tubulin were resuspended in an equal volume of 2× sample buffer corresponding to the soluble fraction volume, followed by sonication and heat denaturation at 95°C for 5 min. Tubulin fractions were subsequently analyzed by SDS-PAGE and Western blot. For cell cycle analysis, treated cells were fixed in 70% ethanol, stained with propidium iodide, and analyzed by flow cytometry (FACSCalibur, BD). Data were analyzed using FlowJo (v10.9.0).

### Cellular uptake of adiposomes

H929 and Raji cells were cultured and adjusted to a density of 1 × 10^6^ cells/mL. Cells were incubated with PBS, fluorescent adiposomes, fluorescent Bio-Ad, or fluorescent BCMA@Ad at 37°C for 1 h. For blocking experiments, BCMA@Ad was preincubated with recombinant BCMA protein prior to incubation with H929 cells. After incubation, suspension cells were collected by centrifugation at 300*g*, washed with PBS, and resuspended for flow cytometry or confocal imaging. HEK293 cells stably expressing LDLR-mCherry were seeded onto confocal dishes and cultured to approximately 70% confluence. Cells were switched to serum-free medium and starved overnight before treatment. Cells were then incubated with fluorescent adiposomes or fluorescent Liver@Ad at 37°C for 1 h. After incubation, cells were washed with PBS, stained with Hoechst for nuclei, and subjected to confocal imaging.

### *In vivo* biodistribution analysis

For lung-targeting analysis, female BALB/c mice were intravenously injected with saline, fluorescent adiposomes, or fluorescent Lung@Ad. Three hours after injection, mice were euthanized and major organs were harvested, rinsed with saline, and imaged using an IVIS system.

For real-time visualization of Liver@Ad uptake, male C57BL/6J mice (8 weeks old) were intravenously injected with 200 μL fluorescent Liver@Ad. Five minutes before imaging, Hoechst 33342 (10 mg/mL) and Evans blue (Bioroyee, #LR1242, 10 mg/mL) were intravenously administered at a volume of 100 μL. Thirty minutes after adiposome administration, mice were anesthetized and the liver was surgically exposed through a small abdominal incision. A liver lobe was gently positioned for imaging using a JP-01 intravital imaging system (Super Vision, Nanjing, China). Images were analyzed using ImageJ software.

### Western blot analysis

Silver staining of proteins was performed as described method previously1. Samples were separated by SDS-PAGE and transferred onto PVDF membranes. Membranes were blocked with 5% non-fat milk and incubated with primary antibodies overnight at 4°C, followed by incubation with HRP-conjugated secondary antibodies. Protein signals were visualized using enhanced chemiluminescence (ECL) reagents and quantified using ImageJ software.

The following primary antibodies were used in this study: β-Tubulin (Abclonal, #A12289, 1:5,000), β-Actin (Abclonal, #AC038, 1:10,000), GAPDH (Abclonal, #AC033, 1:50,000), GFP (Roche, #11814460001, 1:1,000), NRP1 (Abclonal, #A19087, 1:2,000), BCMA (ABMAX, #WPS001, 1:1,000), ApoE (Abclonal, #A0304SP, 1:2,000), and LDLR (Abclonal, #A14996SP, 1:1,000). HRP-conjugated goat anti-rabbit IgG (Abclonal, #AS014, 1:10,000) and goat anti-mouse IgG (Abclonal, #AS003, 1:10,000) were used as secondary antibodies.

### Hemolysis assay

Fresh blood was collected from healthy female BALB/c mice, and red blood cells (RBCs) were isolated by centrifugation at 3,000*g* for 10 min. After washing with PBS, RBCs were resuspended in PBS and incubated with indicated formulations at the final docetaxel concentration of 2.5, 5, 10 μg/mL at 37°C for 2 h. PBS and distilled water were used as negative and positive controls, respectively.

After incubation, samples were centrifuged at 3,000*g* for 10 min, and the absorbance of released hemoglobin in the supernatant was measured at 540 nm using a microplate reader. Hemolysis percentage was calculated according to the following equation:

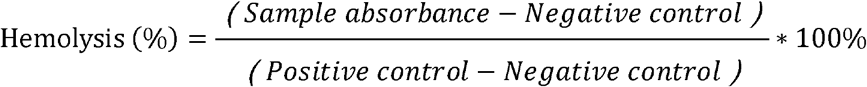

### Tissue distribution of DTX

Male C57BL/6J mice (12-13 weeks old) were intravenously injected with docetaxel at a dose of 20 mg/kg of different formulations. At 1 h post-injection, liver and lung were collected. Tissues were rinsed with saline, blotted dry, weighed, and stored at -80°C until analysis. For docetaxel quantification, tissue samples were homogenized and DTX was extracted with acetonitrile. Paclitaxel was used as an internal standard. The samples were subjected to LC-MS/MS analysis. Standard curves were generated using docetaxel standards prepared in biological matrices. The ratio of analyte to internal standard peak area was used for linear regression, and docetaxel concentrations in tissue samples were calculated accordingly.

### Subcutaneous tumor models

Female BALB/c-nu mice (8 weeks old) were subcutaneously inoculated with MCF-7 cells (2 × 10^6^ cells per mouse) or H226 cells (5 × 10^6^ cells per mouse). Female BALB/c mice were subcutaneously inoculated with 4T1-Luc cells (1.5 × 10^5^ cells per mouse). Tumor size was measured using calipers, and tumor volume was calculated as:

V = L × W^2^ × 0.5

where L and W represent the longest and shortest tumor diameters, respectively. Mice were randomized into treatment groups when tumors reached comparable sizes.

DTX-Inj or DTX-Ad was administered via tail vein injection at indicated docetaxel-equivalent doses (15 mg/kg for MCF-7 and 4T1-Luc models; 20 mg/kg for H226 model), while saline was administered at equal volumes as control. PTX-HSA was included in the MCF-7 model and administered via tail vein injection at 15 mg/kg. Treatments were performed on days 1, 8, and 15 for MCF-7 and H226 models, and on days 1, 7, and 14 for the 4T1-Luc model. Tumor growth and body weight were monitored throughout treatment. Mice were euthanized at predefined experimental endpoints. Tumors and major tissues were collected for downstream analyses.

### Lung tumor colonization model

Female BALB/c mice (8 weeks old) were intravenously injected with 5 × 10^4^ 4T1-Luc cells. On day 4, tumor colonization was assessed by IVIS imaging, and mice were randomized into treatment groups based on comparable bioluminescence intensities. Indicated formulations were administered via tail vein injection at a docetaxel-equivalent dose of 8 mg/kg on days 4, 6, and 8. Tumor progression was monitored by bioluminescence imaging. Mice were euthanized at predefined experimental endpoints.

### Multiple myeloma model

A systemic multiple myeloma model was established by intravenous injection of 1 × 10^7^ U266-Luc cells into female NCG mice (8 weeks old). Twenty-eight days later, tumor burden was assessed by IVIS imaging, and mice were randomized into treatment groups based on comparable luminescence intensities. Mice were treated via tail vein injection with indicated formulations at docetaxel-equivalent doses of 20 mg/kg on days 28, 33, 39, and 46. Tumor progression was monitored by bioluminescence imaging. Mice were euthanized at predefined experimental endpoints.

### Orthotopic hepatocellular carcinoma model

An orthotopic hepatocellular carcinoma model was established using Hepa1-6 cells. Male C57BL/6J mice (8 weeks old) were anesthetized with tribromoethanol, and 2 × 10^6^ Hepa1-6 cells mixed with Matrigel were surgically implanted into the liver. The abdominal incision was closed after implantation, and mice were randomized into treatment groups 3-5 days later. DTX formulations were administered via tail vein injection at a docetaxel-equivalent dose of 15 mg/kg on days 0, 7, and 14. Sorafenib (Aladdin, #S129593) was prepared at 5 mg/mL in Cremophor EL/95% ethanol/H[O (1:1:6, v/v/v) and administered daily by oral gavage at approximately 40 mg/kg. Mice were euthanized at predefined experimental endpoints and tissues were collected for downstream analyses.

### Chemically induced spontaneous hepatocellular carcinoma model

Male C57BL/6J mice were intraperitoneally injected with DEN (40 mg/kg) at 1 week of age. After 6 weeks, mice received intraperitoneal injections of 20% carbon tetrachloride in corn oil (5 μL/g) twice weekly for 12 weeks. Mice were then randomized into treatment groups according to body weight. DTX formulations were administered via tail vein injection at a docetaxel-equivalent dose of 15 mg/kg on days 0, 7, and 14. Saline and blank adiposome controls were administered intravenously at equivalent volumes. Body weight was monitored. Mice were euthanized at predefined experimental endpoints and tissues were collected for downstream analyses.

### Statistical analysis

Data are presented as mean ± SEM unless otherwise indicated. Statistical analyses were performed using GraphPad Prism 9.0.0. Two-tailed Student’s *t* test was used for comparisons between two groups unless otherwise specified. Differences were considered statistically significant at p < 0.05.

**The other method sections were provided in the supplementary files**.

## Reference

[1] Subramanian G, Adams M D, Venter J C, et al. Implications of the human genome for understanding human biology and medicine [J]. JAMA, 2001, 286(18): 2296–307.

[2] Sarkar C, Das B, Rawat V S, et al. Artificial Intelligence and Machine Learning Technology Driven Modern Drug Discovery and Development [J]. International Journal of Molecular Sciences, 2023, 24(3).

[3] Leeson P D, Springthorpe B. The influence of drug-like concepts on decision-making in medicinal chemistry [J]. Nature reviews Drug discovery, 2007, 6(11): 881–90.

[4] Zhang R, Qin X, Kong F, et al. Improving cellular uptake of therapeutic entities through interaction with components of cell membrane [J]. Drug Delivery, 2019, 26(1): 328–42.

[5] Kalepu S, Nekkanti V. Insoluble drug delivery strategies: review of recent advances and business prospects [J]. Acta Pharmaceutica Sinica B, 2015, 5(5): 442–53.

[6] Yang H, Wang Y, Liu W, et al. Genome-wide pan-GPCR cell libraries accelerate drug discovery [J]. Acta Pharmaceutica Sinica B, 2024, 14(10): 4296–311.

[7] Sriram K, Insel P A. G Protein-Coupled Receptors as Targets for Approved Drugs: How Many Targets and How Many Drugs? [J]. Molecular Pharmacology, 2018, 93(4): 251–8.

[8] Mateus A, Gordon L J, Wayne G J, et al. Prediction of intracellular exposure bridges the gap between target- and cell-based drug discovery [J]. Proceedings of the National Academy of Sciences of the United States of America, 2017, 114(30): E6231–E9.

[9] Liu J, Cabral H, Mi P. Nanocarriers address intracellular barriers for efficient drug delivery, overcoming drug resistance, subcellular targeting and controlled release [J]. Advanced Drug Delivery Reviews, 2024, 207: 115239.

[10] Mitchell M J, Billingsley M M, Haley R M, et al. Engineering precision nanoparticles for drug delivery [J]. Nat Rev Drug Discov, 2020, 20(2): 101–24.

[11] Bulbake U, Doppalapudi S, Kommineni N, et al. Liposomal formulations in clinical use: an updated review, Pharmaceutics 9 (2017) 12 [Z]. DOI. 2016

[12] Kumar S, Randhawa J K. High melting lipid based approach for drug delivery: solid lipid nanoparticles [J]. Mater Sci Eng C Mater Biol Appl, 2013, 33(4): 1842–52.

[13] Li B, Yuan Z, Hung H-C, et al. Revealing the Immunogenic Risk of Polymers [J]. Angew Chem Int Ed Engl, 2018, 57(42): 13873–6.

[14] Feingold K R. Introduction to Lipids and Lipoproteins [M]. South Dartmouth (MA): MDText.com, Inc., 2024.

[15] Makoto K, Kazuhisa T, Shengyi L, et al. Minor apolipoproteins contribute to the pleiotropic effects of high-density lipoprotein [J]. Biophysics Reports, 2026, 0(0).

[16] Wang Y, Zhou X-M, Ma X, et al. Construction of Nanodroplet/Adiposome and Artificial Lipid Droplets [J]. ACS Nano, 2016, 10(3): 3312–22.

[17] Al-Jipouri A, Almurisi S H, Al-Japairai K, et al. Liposomes or Extracellular Vesicles: A Comprehensive Comparison of Both Lipid Bilayer Vesicles for Pulmonary Drug Delivery [J]. Polymers, 2023, 15(2).

[18] Fu Y, Ding B, Liu X, et al. Qa-SNARE syntaxin 18 mediates lipid droplet fusion with SNAP23 and SEC22B [J]. Cell Discovery, 2023, 9(1): 115.

[19] Wan N, Hong Z, Parson M A H, et al. Spartin-mediated lipid transfer facilitates lipid droplet turnover [J]. Proceedings of the National Academy of Sciences of the United States of America, 2024, 121(3): e2314093121.

[20] Dias AraúJo A R, Bello A A, Bigay J, et al. Surface tension-driven sorting of human perilipins on lipid droplets [J]. The Journal of Cell Biology, 2024, 223(12).

[21] Zhang C, Liu P. The New Face of the Lipid Droplet: Lipid Droplet Proteins [J]. Proteomics, 2018, 19(10): e1700223.

[22] Zhang C, Yang L, Ding Y, et al. Bacterial lipid droplets bind to DNA via an intermediary protein that enhances survival under stress [J]. Nature Communications, 2017, 8(1): 15979.

[23] Peng H, Xu Q, Zhang T, et al. Molecular determinants for the association of human hormone-sensitive lipase with lipid droplets [J]. Nature Communications, 2025, 16(1): 3497.

[24] Ma X, Zhi Z, Zhang S, et al. Validating an artificial organelle: Studies of lipid droplet-specific proteins on adiposome platform [J]. IScience, 2021, 24(8): 102834.

[25] Cao Z, Zhang Q, Zhou Z, et al. Construction and application of artificial lipoproteins using adiposomes [J]. Journal of Lipid Research, 2023, 64(10).

[26] He W, Wang M, Zhang X, et al. Estrogen Induces LCAT to Maintain Cholesterol Homeostasis and Suppress Hepatocellular Carcinoma Development [J]. Cancer Research, 2024, 84(15): 2417–31.

[27] Engels F K, Mathot R A A, Verweij J. Alternative drug formulations of docetaxel: a review [J]. Anti-cancer Drugs, 2007, 18(2).

[28] Catimel G, Verweij J, Mattijssen V, et al. Docetaxel (Taxotere): an active drug for the treatment of patients with advanced squamous cell carcinoma of the head and neck. EORTC Early Clinical Trials Group [J]. Annals of Oncology : Official Journal of the European Society For Medical Oncology, 1994, 5(6): 533–7.

[29] Valero V, Holmes F A, Walters R S, et al. Phase II trial of docetaxel: a new, highly effective antineoplastic agent in the management of patients with anthracycline-resistant metastatic breast cancer [J]. Journal of Clinical Oncology : Official Journal of the American Society of Clinical Oncology, 1995, 13(12): 2886–94.

[30] Tan Q, Liu X, Fu X, et al. Current development in nanoformulations of docetaxel [J]. Expert Opinion On Drug Delivery, 2012, 9(8): 975–90.

[31] Guéritte-Voegelein F, Guénard D, Lavelle F, et al. Relationships between the structure of taxol analogues and their antimitotic activity [J]. Journal of Medicinal Chemistry, 1991, 34(3): 992–8.

[32] Baker J, Ajani J, Scotté F, et al. Docetaxel-related side effects and their management [J]. European Journal of Oncology Nursing : the Official Journal of European Oncology Nursing Society, 2009, 13(1): 49–59.

[33] Haspel N, Zanuy D, Nussinov R, et al. Binding of a C-End Rule Peptide to the Neuropilin-1 Receptor: A Molecular Modeling Approach [J]. Biochemistry, 2011, 50(10): 1755–62.

[34] Alberici L, Roth L, Sugahara K N, et al. De novo design of a tumor-penetrating peptide [J]. Cancer Res, 2013, 73(2): 804–12.

[35] Liu D, Wang C, Yang J, et al. CRGDK-Functionalized PAMAM-Based Drug-Delivery System with High Permeability [J]. ACS Omega, 2020, 5(16): 9316–23.

[36] Zehmer J K, Bartz R, Bisel B, et al. Targeting sequences of UBXD8 and AAM-B reveal that the ER has a direct role in the emergence and regression of lipid droplets [J]. Journal of Cell Science, 2009, 122(Pt 20): 3694–702.

[37] Wang R, Hu B, Pan Z, et al. Antibody-Drug Conjugates (ADCs): current and future biopharmaceuticals [J]. J Hematol Oncol, 2025, 18(1): 51.

[38] Ochtrop P, Jagtap A P, Felber J G, et al. Expanding the payload scope in antibody-drug conjugates by delivery of hydroxy-containing drugs through self-immolative phosphoramidates [J]. Nature Communications, 2026, 17(1): 759.

[39] Dixit T, Aswini A, Nikam H, et al. Emerging trends in synthesis, characterization, and mechanism of action of antibody-drug and antibody-nanoparticle conjugates [J]. Discov Nano, 2025, 20(1): 139.

[40] Zelepukin I V, Shevchenko K G, Deyev S M. Rediscovery of mononuclear phagocyte system blockade for nanoparticle drug delivery [J]. Nature Communications, 2024, 15(1): 4366.

[41] Zhang L, Zhang N. How nanotechnology can enhance docetaxel therapy [J]. International Journal of Nanomedicine, 2013, 8: 2927–41.

[42] Chaurawal N, Raza K. Nano-interventions for the drug delivery of docetaxel to cancer cells [J]. Health Sciences Review, 2023, 7: 100101.

[43] Eckhoff L, Nielsen M, Moeller S, et al. TAXTOX - a retrospective study regarding the side effects of docetaxel given as part of the adjuvant treatment to patients with primary breast cancer in Denmark from 2007 to 2009 [J]. Acta Oncol, 2011, 50(7): 1075–82.

[44] Zhao P, Astruc D. Docetaxel nanotechnology in anticancer therapy [J]. ChemMedChem, 2012, 7(6): 952–72.

[45] Fahy E, Cotter D, Sud M, et al. Lipid classification, structures and tools [J]. Biochim Biophys Acta, 2011, 1811(11): 637–47.

[46] Kapoor B, Kapoor D, Gautam S, et al. Dietary Polyunsaturated Fatty Acids (PUFAs): Uses and Potential Health Benefits [J]. Curr Nutr Rep, 2021, 10(3): 232–42.

[47] Lunn J, Theobald H E. The health effects of dietary unsaturated fatty acids [J]. Nutrition Bulletin, 2006, 31(3): 178–224.

[48] Yanagita T, Nagao K. Functional lipids and the prevention of the metabolic syndrome [J]. Asia Pac J Clin Nutr, 2008, 17 Suppl 1: 189–91.

[49] Olsen T, Blomhoff R. Retinol, Retinoic Acid, and Retinol-Binding Protein 4 are Differentially Associated with Cardiovascular Disease, Type 2 Diabetes, and Obesity: An Overview of Human Studies [J]. Adv Nutr, 2020, 11(3): 644–66.

[50] Ke X-Y, Zou M, Xu C. Lipid metabolism in tumor-infiltrating T cells: mechanisms and applications [J]. Life Metabolism, 2022, 1(3): 211–23.

[51] Liu P, Ying Y, Zhao Y, et al. Chinese hamster ovary K2 cell lipid droplets appear to be metabolic organelles involved in membrane traffic [J]. J Biol Chem, 2003, 279(5): 3787–92.

[52] Yang L, Ding Y, Chen Y, et al. The proteomics of lipid droplets: structure, dynamics, and functions of the organelle conserved from bacteria to humans [J]. Journal of Lipid Research, 2012, 53(7): 1245–53.

[53] Xu S, Zhang X, Liu P. Lipid droplet proteins and metabolic diseases [J]. Biochim Biophys Acta Mol Basis Dis, 2017, 1864(5 Pt B): 1968–83.

[54] Fu X, Zhang S, Liu P. Co-immunoprecipitation for identifying protein-protein interaction on lipid droplets [J]. Biophys Rep, 2024, 10(2): 102–10.

[55] Teesalu T, Sugahara K N, Kotamraju V R, et al. C-end rule peptides mediate neuropilin-1-dependent cell, vascular, and tissue penetration [J]. Proceedings of the National Academy of Sciences, 2009, 106(38): 16157–62.

[56] Falcon-Rodriguez C I, Osornio-Vargas A R, Sada-Ovalle I, et al. Aeroparticles, Composition, and Lung Diseases [J]. Frontiers in Immunology, 2016, Volume 7 –2016.

[57] Fu Z, Li S, Han S, et al. Antibody drug conjugate: the “biological missile” for targeted cancer therapy [J]. Signal Transduct Target Ther, 2022, 7(1): 93.

[58] Tsuchikama K, Anami Y, Ha S Y Y, et al. Exploring the next generation of antibody-drug conjugates [J]. Nat Rev Clin Oncol, 2024, 21(3): 203–23.

[59] Liu H, Zeng H, Qin X, et al. The Icarian flight of antibody-drug conjugates: target selection amidst complexity and tackling adverse impacts [J]. Protein Cell, 2025, 16(7): 532–56.

[60] Marques A C, Costa P J, Velho S, et al. Functionalizing nanoparticles with cancer-targeting antibodies: A comparison of strategies [J]. J Control Release, 2020, 320: 180–200.

[61] Kalra A, Yetiskul E, Wehrle C J, et al. Physiology, Liver [M]. Treasure Island (FL): StatPearls Publishing, 2023.

[62] Tsoi K M, Macparland S A, Ma X-Z, et al. Mechanism of hard-nanomaterial clearance by the liver [J]. Nature Materials, 2016, 15(11): 1212–21.

[63] Ngo W, Ahmed S, Blackadar C, et al. Why nanoparticles prefer liver macrophage cell uptake in vivo [J]. Advanced Drug Delivery Reviews, 2022, 185: 114238.

[64] Harkins L, Vilarinho S, Saltzman W M. Targeting Polymeric Nanoparticles to Specific Cell Populations in the Liver [J]. Biochemistry, 2025, 64(8): 1685–97.

[65] Peng M, Fang F, Wang B. Nanoparticle technologies for liver targeting and their applications in liver diseases [J]. Front Bioeng Biotechnol, 2025, 13: 1661872.

[66] Cao Z, Lei L, Zhou Z, et al. Apolipoprotein A-IV and its derived peptide, T55™121, improve glycemic control and increase energy expenditure [J]. Life Metabolism, 2024, 3(4): loae010.

[67] Jeon H, Blacklow S C. Structure and physiologic function of the low-density lipoprotein receptor [J]. Annu Rev Biochem, 2005, 74(1): 535–62.

[68] Huang Y, Mahley R W. Apolipoprotein E: structure and function in lipid metabolism, neurobiology, and Alzheimer’s diseases [J]. Neurobiology of disease, 2014, 72: 3–12.

