## Supplementary data-20260608 for "Nature-Inspired Nanoparticle Adiposomes Enable Targeted Delivery of Hydrophobic Drug for Anti-Cancer Treatment"

**Supplementary methods**

***Docetaxel solubility screening in neutral lipids***

Docetaxel was mixed with different neutral lipids, including corn oil, fish oil, tricaprylin, or mixed neutral lipids, at indicated concentrations. After vortexing and incubation, samples were centrifuged at 20,000*g* for 5 min to remove undissolved drug when necessary. The supernatant or lipid phase was collected and analyzed by HPLC to determine docetaxel solubility. Nile Red (Thermo Fisher Scientific, #N1142) staining was used to visualize adiposomes during formulation screening.

***Preparation of fluorescence-labeled adiposomes***

Fluorescence-labeled adiposomes were prepared using different labeling strategies depending on experimental purposes.

For intrinsic fluorescence labeling, adiposomes were prepared by incorporating 40 μg TopFluor glycerol (Avanti Polar Lipids, #810271) into 200 μL neutral lipids prior to adiposome assembly. Fluorescent adiposomes were prepared using the same procedure as blank adiposomes.

For neutral lipid-specific staining, adiposomes were stained with Nile Red, LipidTOX Red or LipidTOX Green (Thermo Fisher Scientific, #H34476 and #H34475) prior to imaging. Stained adiposomes, including those incubated with GFP-avidin or ApoE-mApple, were imaged using fluorescence microscopy (ZEISS Axio Imager M2) or confocal laser scanning microscopy (OLYMPUS FV3000RS).

***Dynamic light scattering and zeta-potential analysis***

Particle size distribution, PDI, and zeta-potential of adiposomes were measured using the BeNano 90 Zeta analyzer (Bettersize Instruments, Dandong, China). For each measurement, 5 μL of adiposome suspension was diluted into 995 μL of ultrapure water, gently mixed, and transferred into a cuvette or zeta cell for the measurement according to the manufacturer’s instructions.

***Dialysis-based stability, release, and particle size analysis***

Dialysis experiments were performed using a dialysis device (Thermo Fisher Scientific, #88402). Briefly, 500 μL of DTX-Ad was loaded into the dialysis cup, and phosphate-buffered saline (PBS) was used as the external medium. Dialysis was conducted at room temperature with horizontal shaking at 80 rpm, and the external PBS was replaced at regular intervals. At the indicated time points, samples were collected from the dialysis device and subjected to HPLC (Agilent, 1260 Infinity) analysis for docetaxel quantification. In parallel, aliquots were taken for measurement of particle size and PDI during the dialysis process.

***Thin-layer chromatography***

Total lipids of adiposome-based formulations were extracted using a mixture of aqueous phase, methanol, and chloroform (1:1:2, v/v/v). After phase separation, the chloroform layer was collected and dried under a gentle stream of nitrogen. The dried lipids were redissolved in 50 μL chloroform and applied onto silica gel plates. For separation of neutral lipids, plates were developed in hexane/diethyl ether/acetic acid (80:20:1, v/v/v). For separation of phospholipids, plates were developed in chloroform/methanol/acetic acid/water (75:13:9:3, v/v/v/v). Lipid bands were visualized under saturated iodine vapor, photographed, and quantified by grayscale analysis using ImageJ software.

***Cell viability assay***

Cells were seeded in 96-well plates at approximately 2,000-5,000 cells per well depending on proliferation rate. Cells were then treated with indicated formulations at the indicated concentration and duration. After treatment, cell viability was determined using CCK-8 reagent (Lablead, #CK001) according to the manufacturer’s instructions. Cell viability was normalized to the untreated control group.

***Confocal imaging of microtubule organization***

Sterilized glass coverslips were placed into 12-well plates, and MCF-7 cells were seeded and cultured to approximately 70% confluence. Cells were treated with medium, DTX-Inj, or DTX-Ad at a docetaxel-equivalent concentration of 50 ng/mL for 24 h. Cells were then washed with PBS, fixed with paraformaldehyde, stained with Hoechst (Beyotime, #C1022) for nuclei and Tubulin-Tracker Red (Beyotime, #C1050) for microtubules, and imaged using an OLYMPUS FV3000RS confocal microscope.

***Proteomic analysis***

Hepa1-6 cells were treated with control, DTX-Inj, or DTX-Ad at a docetaxel-equivalent concentration of 100 ng/mL for 12 h. After treatment, cells were washed twice with PBS. Cells were then lysed using 6 M urea in 0.1 M triethylammonium bicarbonate (TEAB, Sigma, #T7408), rapidly frozen in liquid nitrogen, and stored at -80°C until analysis. Proteomic analysis was performed by a commercial service provider (NovoMagic, China) using an Orbitrap Astral mass spectrometer. Protein digestion, LC-MS/MS acquisition, and primary data processing were conducted according to standard workflows. Differentially expressed proteins were identified using thresholds of P < 0.05 and |log2Fold-change| > 0.58. Functional enrichment analyses were performed using Gene Ontology and pathway databases. Heatmaps, volcano plots, and enrichment analyses were visualized using standard bioinformatic pipelines.

***In vivo imaging and analysis***

Bioluminescence and fluorescence imaging were performed using an *In Vivo* Imaging System (IVIS, PerkinElmer, Lumina III). D-luciferin (Lablead, #D22505) was administered intraperitoneally at 0.15 mg/g at 10 min before imaging. Bioluminescence and fluorescence signals were quantified using Living Image software (v4.3.1).

***Histological, hematological, and serum biochemical analyses***

Major tissues were fixed, embedded, sectioned, and subjected to hematoxylin and eosin staining for histological evaluation. Blood samples were analyzed for hematological indices and serum biochemical markers using standard clinical assays. Some histological staining and blood routine analyses were performed by qualified institutional or commercial service platforms.

Supplementary Figure S1

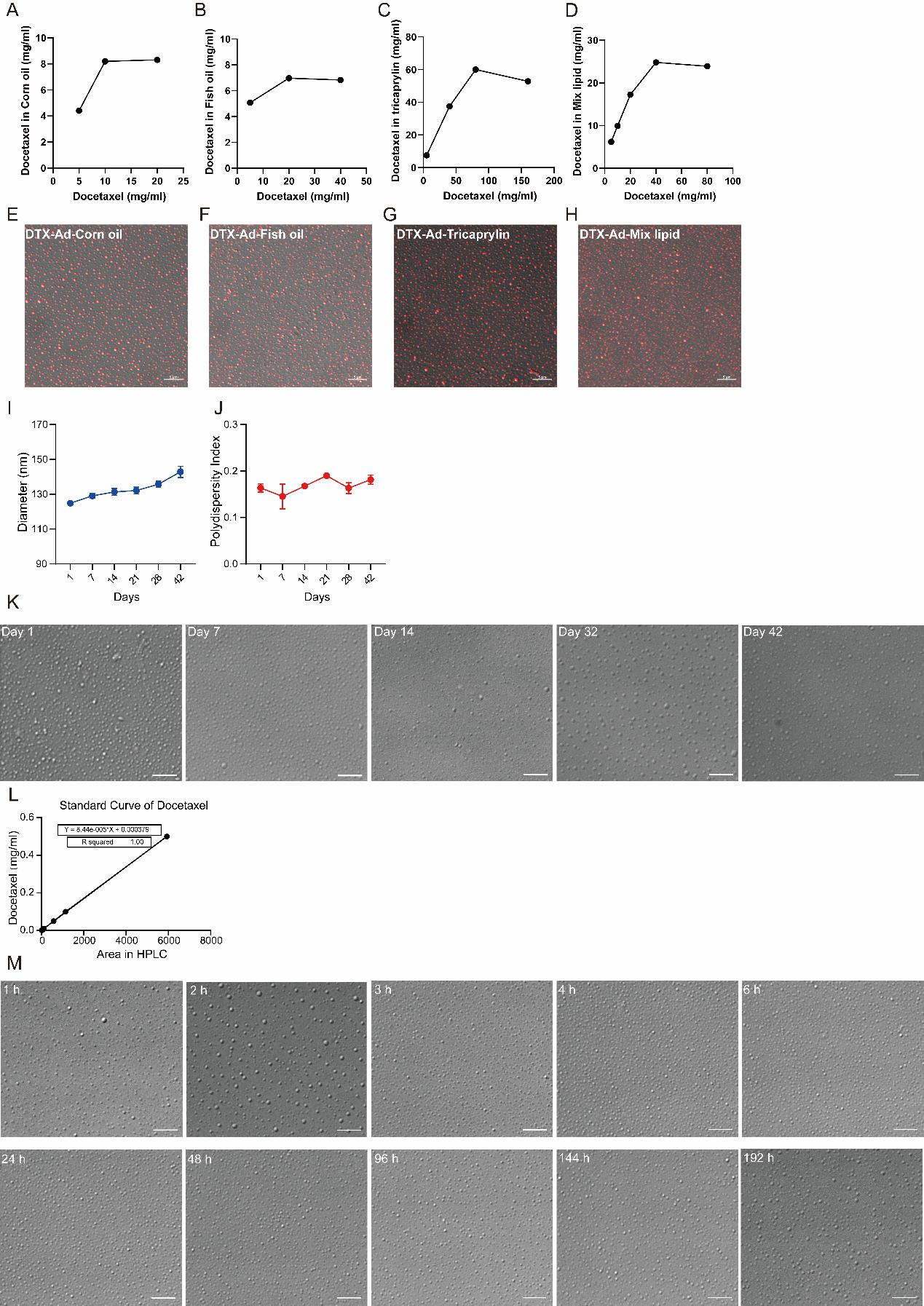

Supplementary Figure S2

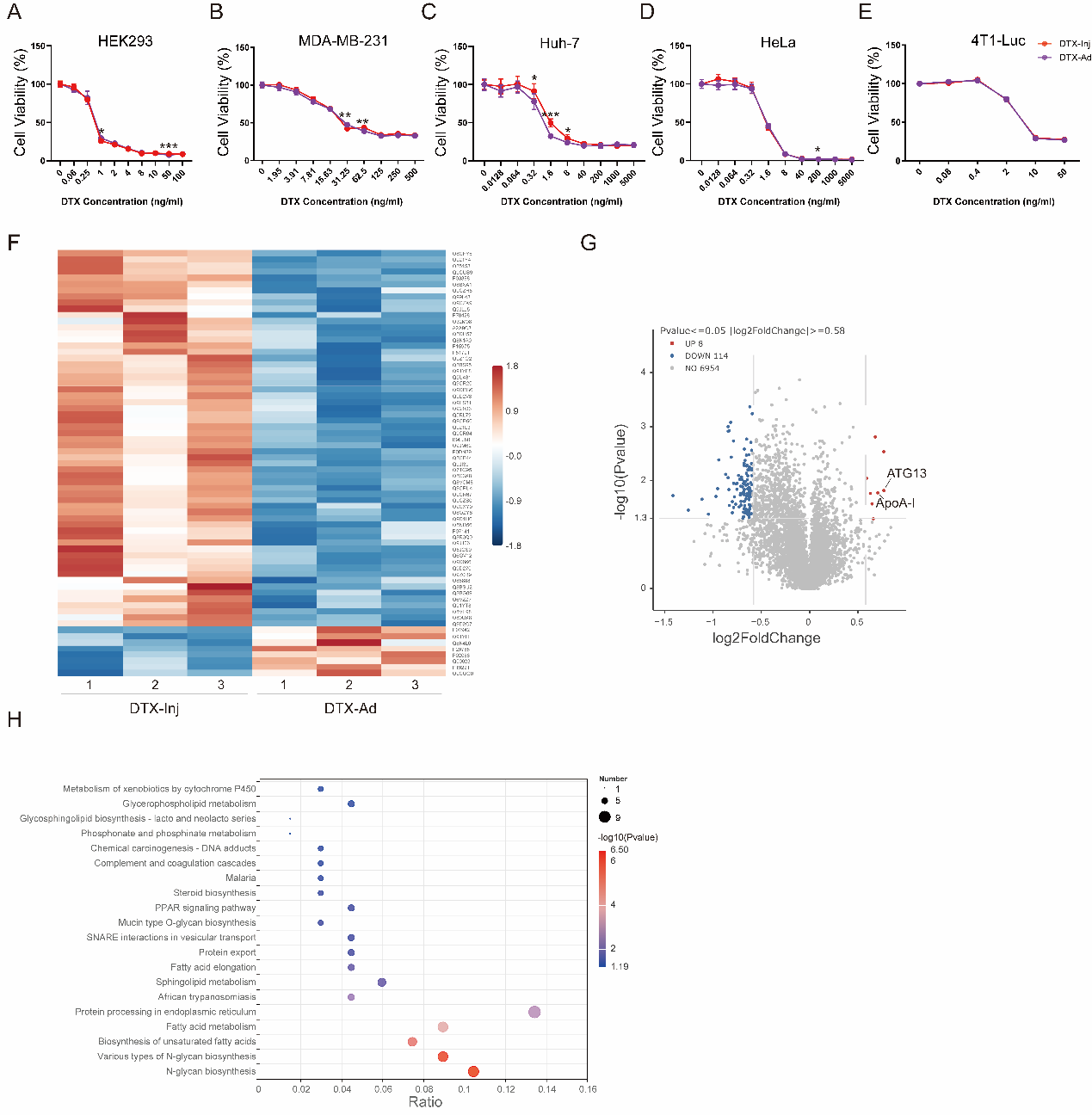

Supplementary Figure S3

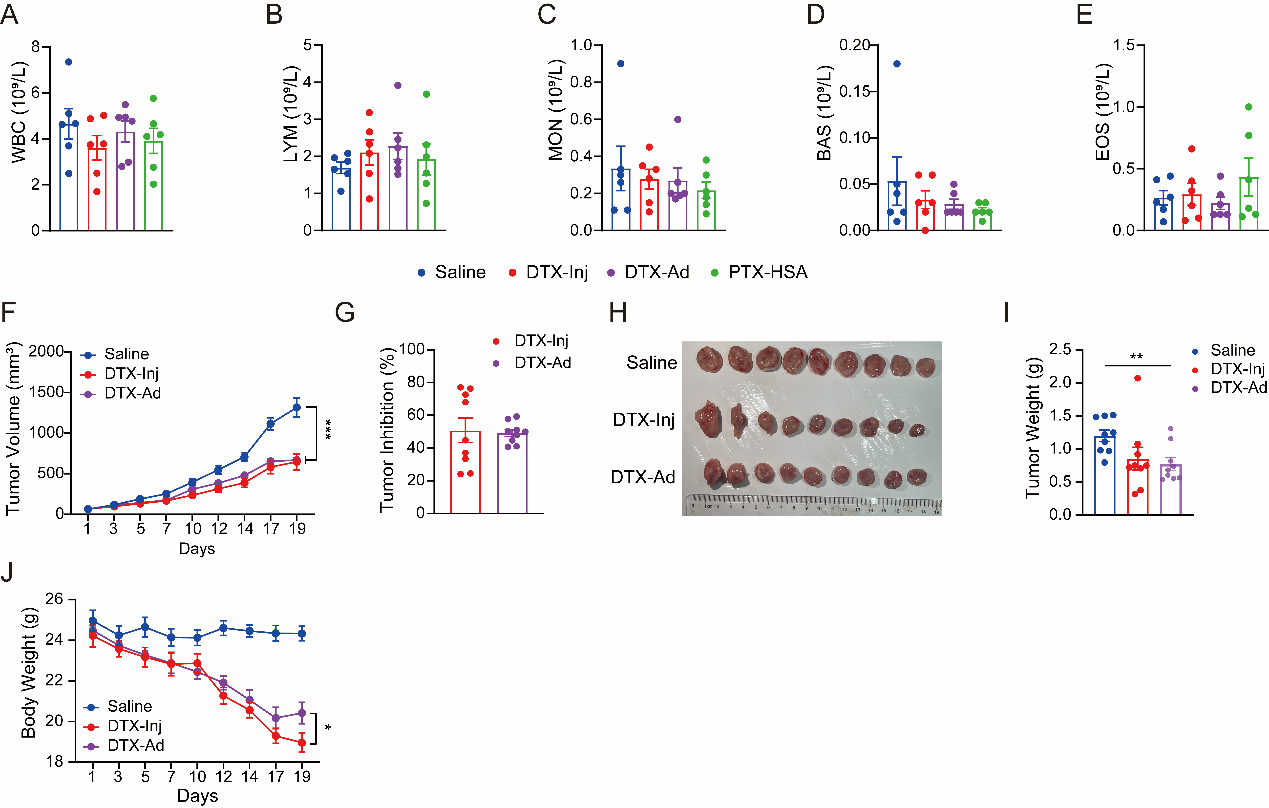

Supplementary Figure S4

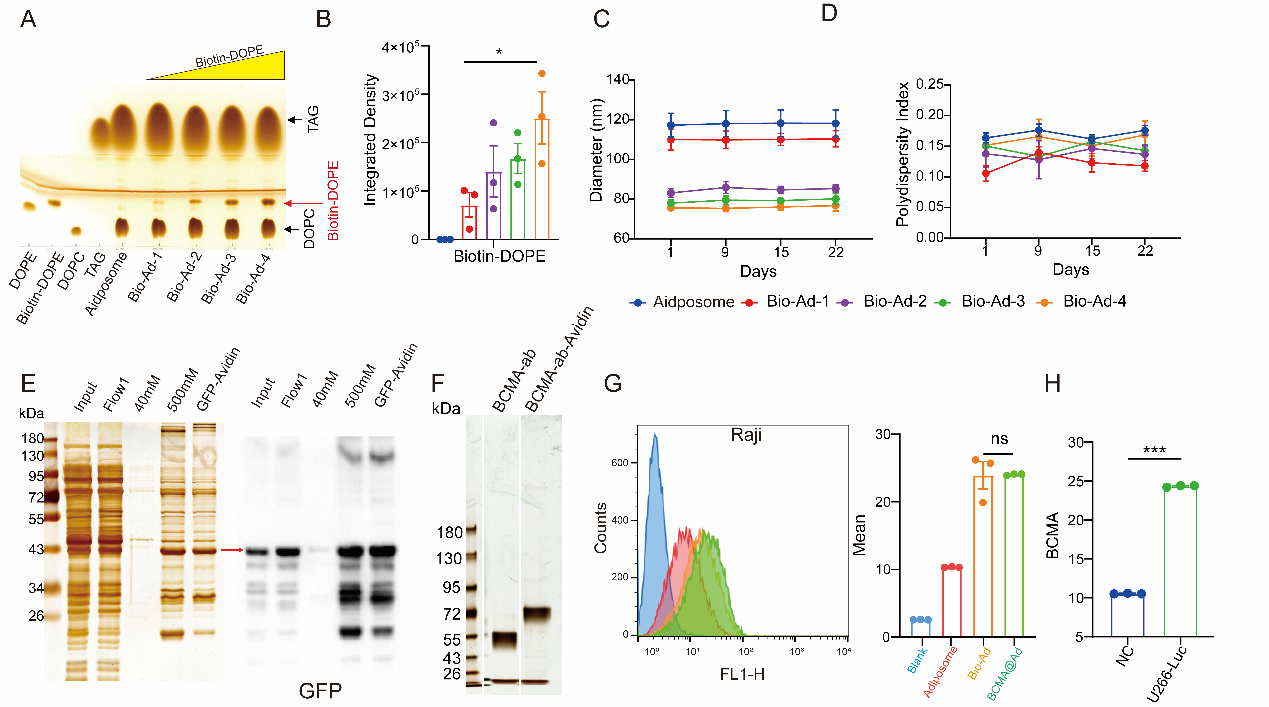

Supplementary Figure S5

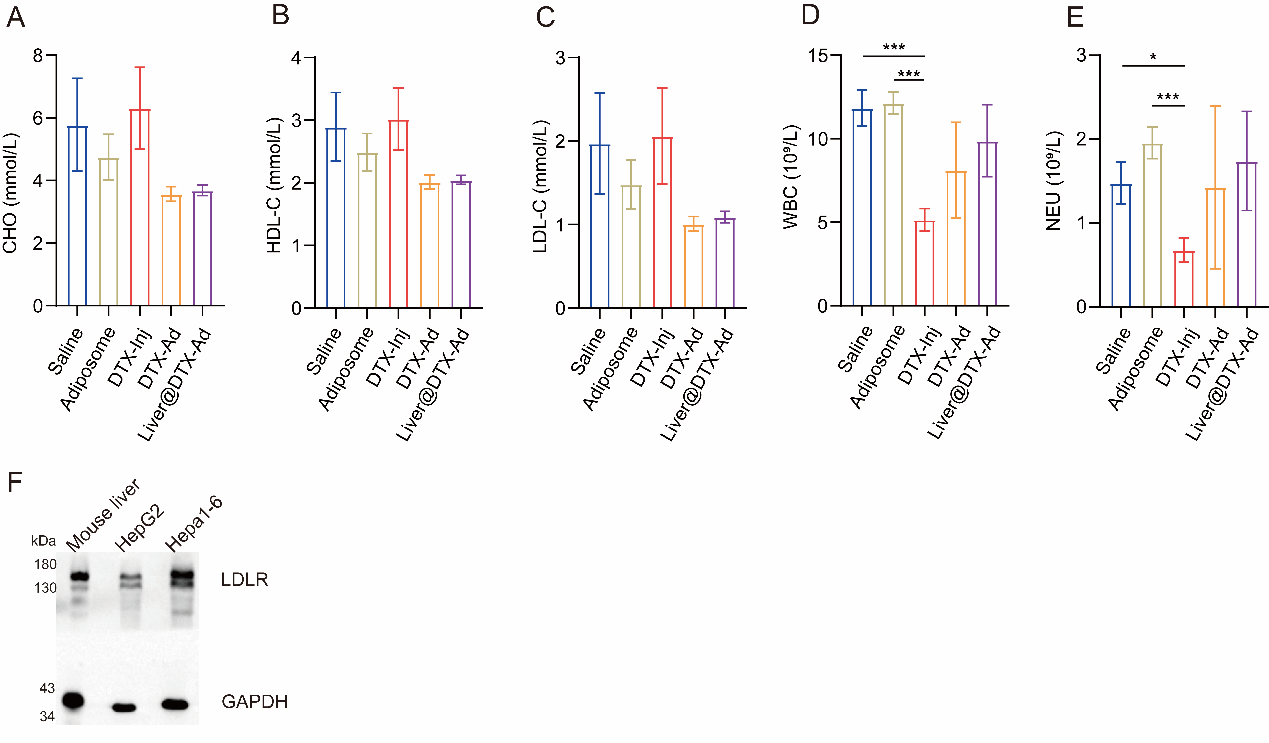

**Figure legend**

**Supplementary Figure S1 Optimization of lipid composition and stability characterization of adiposomes.**
(A-D) Solubility of DTX in corn oil, fish oil, tricaprylin and mixed lipids (n = 1 or 2).
(E-H) Fluorescence images of DTX-Ad prepared with different neutral lipid compositions. Scale bars: 5 μm.

(I-K) Particle size, PDI and images of blank adiposomes during storage at 4 °C (n = 3). Scale bar: 5 μm.
(L) HPLC standard curve of DTX.
(M) Morphological observation of DTX-Ad during dialysis in PBS. Scale bar: 5 μm.

Data are presented as mean values or mean ± SEM.

**Supplementary Figure S2 Extended cytotoxicity and proteomic analysis of DTX-Ad.**
(A-E) Cell viability of HEK293, MDA-MB-231, Huh-7, HeLa and 4T1-Luc cells treated with DTX-Inj, DTX-Ad or blank adiposomes at the indicated concentrations, measured by CCK-8 assay.

(F)  Heatmap of differentially expressed proteins in Hepa1-6 cells treated with DTX-Inj or DTX-Ad for 12 h at a docetaxel-equivalent concentration of 100 ng/mL (n = 3).

(G) Volcano plot of differentially expressed proteins, with selected proteins indicated.
(H) KEGG pathway enrichment analysis of differentially expressed proteins.

Data are presented as mean ± SEM. Statistical differences were calculated using two-tailed Student’s *t* test. ****P*<0.001, ***P*<0.01, **P*<0.05.

**Supplementary Figure S3 Safety evaluation and validation of DTX-Ad in additional tumor models.**
**(A-E)** Hematological analysis of white blood cells (WBC), lymphocytes (LYM), monocytes (MON), basophils (BAS), and eosinophils (EOS) in the indicated groups from the MCF-7 xenograft model at the endpoint.

**(F)** Tumor growth curves in the 4T1 syngeneic model established by subcutaneous inoculation of 1.5 × 10^5^ 4T1-Luc cells into BALB/c mice, followed by treatment as indicated.
**(G)** Tumor inhibition rates in the 4T1 model.
**(H)** Representative images of excised tumors from the indicated groups in the 4T1 model.
**(I)** Tumor weights in the 4T1 model at the endpoint.
**(J)** Body weight change in 4T1 tumor-bearing mice during treatment.

Data are presented as mean ± SEM. Statistical differences were calculated using two-tailed Student’s *t* test. ****P*<0.001, ***P*<0.01, **P*<0.05.

**Supplementary Figure S4 Construction and validation of biotin-avidin-mediated antibody functionalization system.**
**(A-B)** TLC analysis and quantification of Biotin-DOPE incorporation into adiposomes (n = 3).
**(C-D)** Particle size and PDI of adiposomes prepared with different Biotin-DOPE incorporation ratios during storage, measured by DLS (n = 3).
**(E)** Silver staining and Western blot analysis of GFP-avidin expression and purification.
**(F)** Silver staining analysis of recombinant BCMA-ab-Avidin fusion protein.
**(G)** Flow cytometry analysis of BCMA@Ad uptake in BCMA-negative Raji cells (n = 3).
**(H)** Western blot analysis of BCMA expression in U266-Luc cells used for the *in vivo* study (n = 3).

Data are presented as mean ± SEM. Statistical differences were calculated using two-tailed Student’s *t* test. ****P*<0.001, **P*<0.05.

**Supplementary Figure S5 Safety evaluation and LDLR expression analysis of ApoE-functionalized adiposomes.**
**(A-C)** Serum lipid profiles including total cholesterol (CHO), HDL-C, and LDL-C.
**(D-E)** Hematological analysis of Liver@DTX-Ad and other groups.

**(F)** Western blot analysis of LDLR expression in mouse liver and hepatoma cell lines.

Data are presented as mean ± SEM. Statistical differences were calculated using two-tailed Student’s *t* test. ****P*<0.001, **P*<0.05.
